# Genome-wide CRISPR interference screen identifies Clip2 as a novel regulator of osteocyte maturation and morphology

**DOI:** 10.1101/2025.10.23.683957

**Authors:** Courtney M. Mazur, Parthena E. Kotsalidis, Majd George, Tom Whalley, Tadatoshi Sato, John G. Doench, Lauren E. Surface, Marc N. Wein

**Affiliations:** Endocrine Unit, Massachusetts General Hospital, Harvard Medical School, Boston, MA, USA; Wellcome Centre for Cell-Matrix Research, Faculty of Biology, Medicine and Health, University of Manchester, Manchester, UK; Department of Medicine, Division of Rheumatology, University of Massachusetts Chan Medical School, Worcester, MA, USA; Broad Institute of MIT and Harvard, Cambridge, MA, USA; University of Michigan School of Dentistry, Ann Arbor, MI, USA

**Keywords:** genome-wide CRISPR screen, osteocyte, cytoskeleton

## Abstract

Osteocytes play critical roles in bone, making them attractive targets for therapeutics to improve bone mass and strength. The genes driving osteocyte maturation and function are not fully understood. Here we aimed to identify novel genes responsible for osteocyte differentiation and dendrite development by performing a genome-wide CRISPR-interference (CRISPRi) screen in the Ocy454 osteocyte-like cell line. We identify CD61 (integrin β3) as a marker of osteocyte maturation: surface CD61 expression increases during osteocyte maturation, and CD61^high^ cells express higher levels of osteocyte marker genes. We then developed a flow cytometry-based assay to quantify surface CD61 protein levels as a phenotypic endpoint for functional genomic screening. In a genome-wide screen, we identified Clip2, which encodes a microtubule binding protein, as one of dozens of genes necessary for CD61 expression. Clip2 inhibition decreased surface CD61 expression, reduced expression of osteocyte-specific genes *Dmp1* and *Sost*, and impaired dendrite morphology *in vitro*. Together, these results highlight the utility of surface CD61 as a marker of osteocyte maturity and identify a role of the microtubule cytoskeleton for osteocyte differentiation, form, and function.

## Introduction

Osteocytes play critical roles in local mineral remodeling, mechanosensation, bone cell coordination, and endocrine signaling [1], [2]. Ablation of osteocytes leads to porous, fragile bone [3]. As osteocytes are post-mitotic, long-lived cells, proper differentiation and morphogenesis have long-term effects on osteocyte function and bone strength. Osteocyte differentiation involves transcriptional and morphologic changes as osteoblasts embed into bone. Although marker genes expressed at different stages of osteoblast-to-osteocyte differentiation have been described [4], [5], [6], the genes that drive osteocyte differentiation and mature morphology remain largely unknown.

Key among the defining characteristics of osteocytes are the numerous dendritic projections extending from each cell body. Dendrites extend through a network of canals in bone, allowing connection to nearby cells via gap junctions [7], [8]. They also massively increase the surface area of osteocytes and house mechanosensitive structures that respond to bone loading [9], [10]. Previously, osteocyte dendrites have been described primarily as actin-rich and actin-dependent structures, with short-term depolymerization of microtubules having little effect on dendrite morphology [11], [12]. However, microtubules contribute to cytoskeletal tension and mechanosensitivity in osteocytes [13], [14], [15], [16], and the role of tubulins and intermediate filament proteins in the extension and long-term maintenance of osteocyte dendrites has not been explored. This leaves open questions regarding the roles of cytoskeletal proteins and their binding partners in establishing osteocyte dendrites and maintaining their functional phenotypes.

Recent work has aimed to define an osteocyte-specific transcriptome that highlights the differences between osteocytes and other cells in the osteoblast lineage [5], [6], [17]. However, the transcriptional changes that drive osteocyte differentiation and acquisition of their dendritic morphology have not been studied at the genome-wide level. Genome-wide clustered regularly interspaced short palindromic repeats (CRISPR)-mediated screens are a high-throughput strategy to measure the effects of individual genetic perturbations on cell viability or phenotype [18]. Optimized and modified Cas9 enzymes now allow for gene ablation through double strand breaks, transcriptional inhibition and activation, base editing, and cleavage or localization of mRNA [19], [20], [21], [22], [23]. CRISPR-based screens have been rapidly adopted to uncover genes controlling drug resistance, cell proliferation & maturation, intracellular protein trafficking & secretion, and cell shape [22], [24], [25], [26], [27], [28], [29], [30].

Here, we sought to identify novel genes responsible for osteocyte differentiation and dendrite development by performing a genome-wide CRISPR-interference (CRISPRi) screen in the Ocy454 osteocyte-like cell line. We developed a flow cytometry-based assay to quantify surface CD61 as a marker of osteocyte maturation and validated our screening results through quantification of osteocyte-specific gene expression and dendrite morphology *in vitro.* Together these results highlight the utility of surface CD61 as a marker of osteocyte maturity and identify a role of the microtubule cytoskeleton in osteocyte differentiation, form, and function.

## Results

### CD61 is a marker of osteocyte maturity

First, in order to perform a flow cytometry-based phenotypic CRISPR screen, we aimed to identify a surface marker to distinguish the differentiation state of osteocytes. We interrogated gene expression data from two different *in vitro* differentiation assays which capture, to some degree, the osteoblast-to-osteocyte transition: Ocy454 cells cultured at 37°C for 14 days versus 1 day and MC3T3-E1 cells cultured in 3D collagen gels compared to growth in 2D on standard polystyrene tissue culture dishes. Ocy454 osteocyte-like cells are conditionally immortalized due to constitutive expression of the thermosensitive large T antigen. Thus, these cells divide at 33°C and stop dividing and up-regulate osteocyte-like gene expression patterns over time at 37°C (**Figure 1A, Supplemental Figure 1A-D, Supplemental Data 1**) [31]. 3D culture of MC3T3-E1 osteoblastic cells induces growth of long dendrite-like cytoskeletal processes and an osteocyte-like transcriptome [5]. Significant upregulation of osteocyte differentiation markers *Sp7*, *Dmp1*, and *Phex* was noted in both systems, and *Sost* was significantly upregulated during Ocy454 differentiation (**Figure 1A-B**).

**Figure 1:**
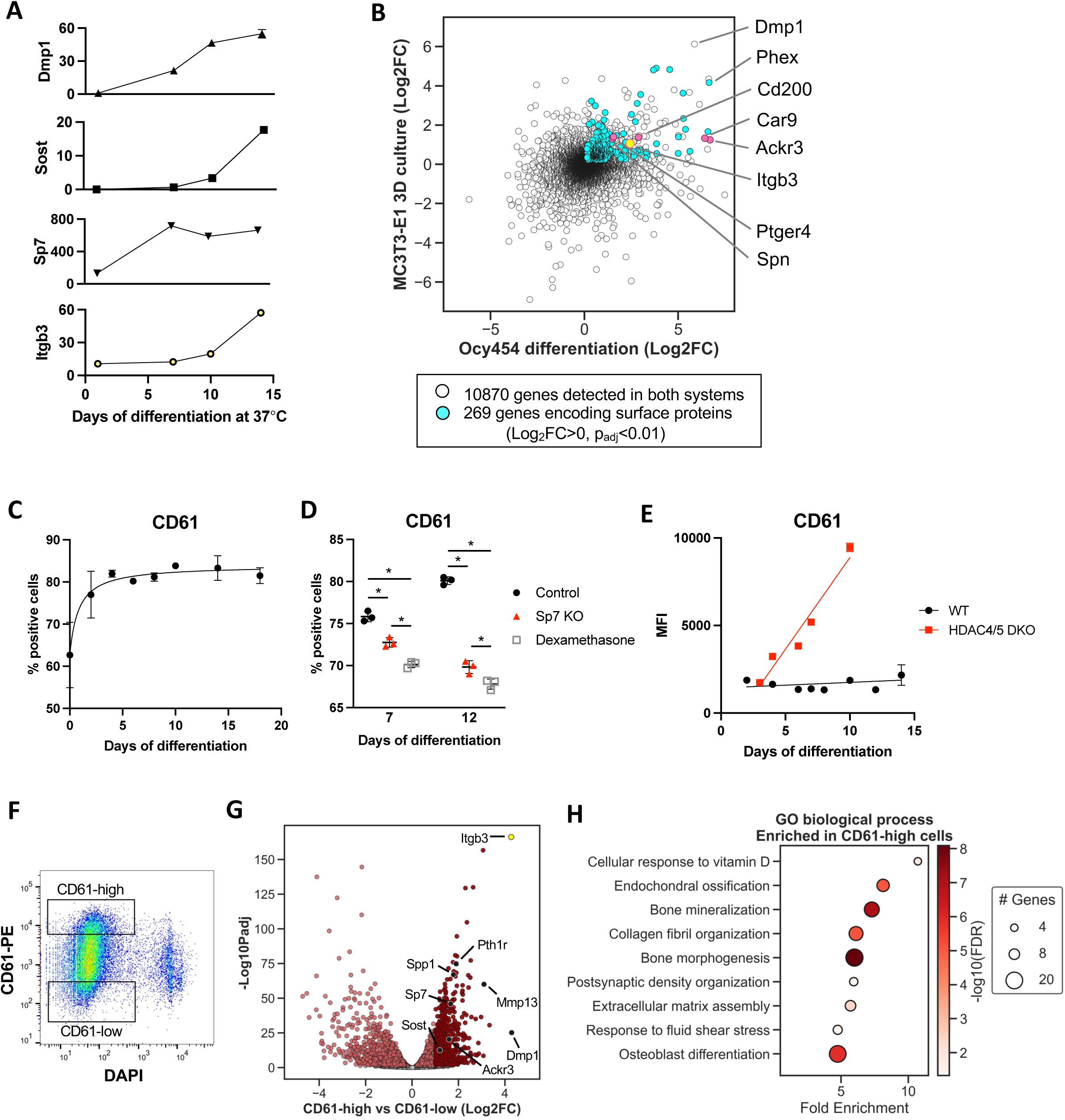
Identification of CD61 as a marker of osteocyte differentiation. A) Counts per million for select osteoblast-osteocyte lineage marker genes from day 1 to day 14 of Ocy454 differentiation in 2D culture by RNA-sequencing. Error bars show standard deviation for two biologic replicates at each time point. B) Scatter plot of genes expressed in Ocy454 cell differentiation (x-axis: Day 14 vs Day 1) and MC3T3-E1 cell 3D vs 2D culture (y-axis). Genes upregulated in both datasets that encode surface proteins are plotted in cyan in the upper right quadrant. Genes encoding candidate flow cytometry markers are plotted in magenta. Itgb3 (plotted in gold, encodes CD61) was ultimately selected. C) Flow cytometry results show percent of cells labeled with CD61-PE antibody compared to isotype control at a range of differentiation timepoints for wild-type Ocy454 cells. Error bars show standard deviation for all replicates at each time point. D) Flow cytometry results show percent of cells labeled with CD61-PE antibody compared to isotype control for wild-type Ocy454 cells, Sp7 knockdown Ocy454 cells, and wild-type Ocy454 cells treated with 1 μM dexamethasone for 24 hours. Error bars show standard deviation for all replicates at each time point. *p<0.01 for all pairwise comparisons, Sidak’s multiple comparisons tests. E) Flow cytometry results show median fluorescence intensity (MFI) of CD61-PE in HDAC4/HDAC5 double knockout Ocy454 cells compared to wild-type Ocy454 cells at a range of differentiation timepoints. Error bars show standard deviation for all replicates of each cell type at each time point. Lines indicate simple linear regression for each cell type, p<0.0001 between genotypes. F) Flow cytometry gates show the DAPI-negative (live) cells with the 10% highest expression of CD61 (CD61-high) and 10% lowest expression of CD61 (CD61-low), collected by FACS for RNA-sequencing after 17 days of differentiation at 37°C. G) Volcano plot of 12,088 genes detected in all samples. 564 genes are significantly enriched in CD61-high cells with Log_2_FC>1 and p_adj_<0.01, n=3 per group. H) Top gene ontology terms enriched in 564 genes upregulated in CD61-high cells.

From these RNA-sequencing datasets, we identified candidate cell surface proteins encoded by genes that were upregulated in both differentiation systems (*Spn*: CD43, *Ackr3*: CXCR7, *Car9*: CA-IX, *Cd200*: CD200, *Ptger4*: PTGER4, *Itgb3*: CD61) and measured their expression on Ocy454 cells by flow cytometry (**Supp Figure 2A-C**). An ideal marker would be strongly upregulated over time at 37°C in Ocy454 cells, detected at high levels on a large percentage of differentiated cells, convenient to use (detected with a commercially-available fluorescently-conjugated antibody), and physiologically relevant.

Of the candidates tested, integrin beta-3 subunit (*Itgb3,* CD61) best met these criteria (**Figure 1C, Supp Figure 2B**). The choice of CD61 as a differentiation marker is bolstered by its physiological relevance in osteocytes. Integrin beta-3 protein is found along osteocyte dendrites *in vivo* [32], [33], and its expression can be modulated by bone-relevant stimuli. Dexamethasone treatment, a clinically relevant pharmacologic insult that reduces osteocyte viability [16], [34], reduces CD61 expression on Ocy454 cells [35] (**Figure 1D**). Furthermore, Ocy454 cells lacking Sp7, a transcription factor needed for dendrite development [5], express less CD61 than WT cells (**Figure 1D**). In contrast, Ocy454 cells lacking HDAC4 and HDAC5, which show accelerated differentiation with high expression of *Sost* [36], have increased CD61 expression (**Figure 1E**).

Next, the physiologic significance of surface CD61 expression was explored by performing RNA-sequencing of CD61^high^ versus CD61^low^ Ocy454 cells after 16 days of differentiation at 37°C (**Figure 1F**). As expected, CD61^high^ cells show increased *Itgb3* expression versus CD61^low^ cells (**Figure 1G**). Gene ontology analysis reveals that CD61^high^ cells are enriched for genes related to ossification, ECM organization, and cell adhesion, such as *Dmp1, Sost, Spp1, Mmp13, Sp7, Pth1r,* and *Ackr3 (***Figure 1G-H, Supplemental Figure 3, Supplemental Data 2**). Together, these results support the choice of surface CD61 as a marker of osteocyte maturity for subsequent genome wide screening.

### Design of a genome-wide CRISPRi screen in Ocy454 cells

We aimed to identify genes that, when inhibited, interfere with Ocy454 cell maturation, detected as reduced CD61 surface expression. Due to concerns that double stranded DNA breaks might affect cell proliferation and differentiation in this long term assay [37], we chose to perform a CRISPR-interference (CRISPRi) screen. We transduced Ocy454 cells with a BFP-expressing lentivirus expressing nuclease-deficient Cas9 (dCas9) fused with transcriptional repressor KRAB [21], [38], and sorted twice for BFP^high^ cells. Similar to parental cells, the resulting “dCas9-Ocy454 cells” upregulated CD61 between 0 and 10 days of differentiation (**Figure 2A-B**). To demonstrate functionality of the dCas9 system, we introduced three unique sgRNAs targeting the *Itgb3* promoter. After 11 days of differentiation, all three sgRNAs reduced the percentage of CD61-positive cells from approximately 70% to 10% (**Figure 2C-D**).

**Figure 2:**
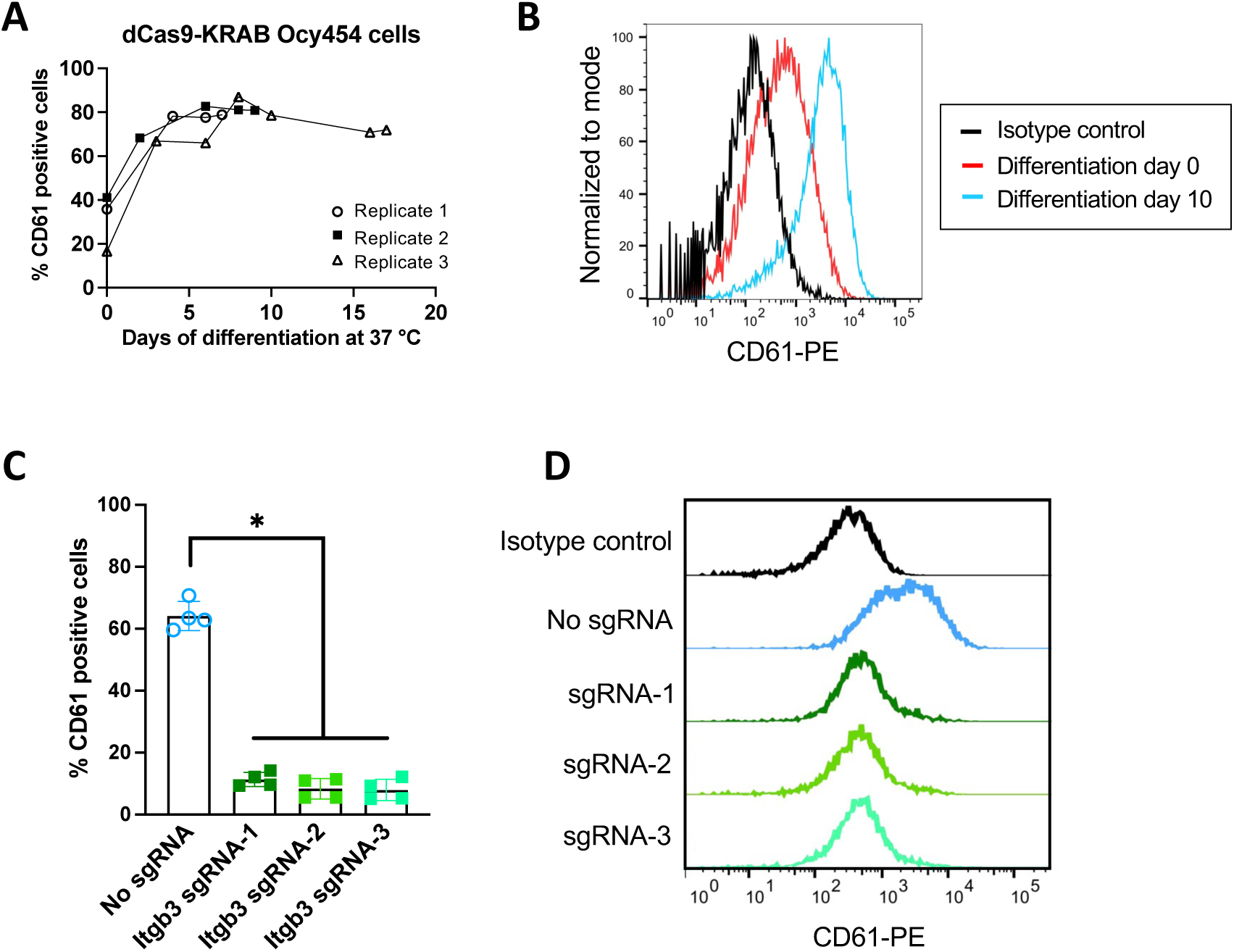
CRISPRi-mediated CD61 silencing in Ocy454 cells. A) Flow cytometry results show percentage of dCas9-KRAB-expressing Ocy454 cells labeled with CD61-PE antibody compared to isotype control at a range of differentiation timepoints. Lines connect matched subcultures of cells followed over time. Error bars show standard deviation for technical replicates in each test. B) Histograms show intensity of CD61-PE fluorescence compared to isotype control in Ocy454 cells differentiated for the indicated times. C) Flow cytometry results show percent of dCas9-KRAB-expressing Ocy454 cells labeled with CD61-PE antibody compared to isotype control in cells with no sgRNA and cells expressing sgRNAs targeting the Itgb3 promoter at differentiation day 11. Each point represents one biological replicate. Error bars show standard deviation. *p<0.05. D) Histograms show intensity of CD61-PE fluorescence compared to isotype control in cells expressing sgRNAs targeting the Itgb3 promoter.

The percentage of cells labeled and labeling intensity of CD61 were insensitive to the volume of antibody used between 0.25-2 µL per 1,000,000 cells (**Supp Figure 4A**). As Ocy454 cells differentiate in 2D culture, they produce extracellular matrix that must be digested to dissociate cells into a single cell suspension for flow cytometry. We compared multiple cell dissociation enzymes and found that trypsin resulted in lower median fluorescence intensity of CD61-PE than the other enzymes tested (**Supp Figure 4B**). However, while all enzymes tested were sufficient to liberate undifferentiated osteocytes, highly concentrated Trypsin-EDTA yielded the most live differentiated cells for analysis (**Supp Figure 4C**). This made carefully-timed application of Trypsin-EDTA the preferred method for liberation of matrix-embedded Ocy454 cells for flow cytometry analysis.

### CRISPRi screen

For screening, we initially aimed to express each of 66,000 sgRNAs in the pooled library in 200 cells (200x representation). To avoid cells simultaneously infected with multiple sgRNA-expressing lentiviruses, we performed titer calculations to achieve lentiviral infection efficiency between 30-50% (**Supp Fig 5A-C**). After pooled library infection and puromycin selection, the heterogeneous population of sgRNA-expressing dCas9-Ocy454 cells proliferated at 33°C until confluent, then differentiated for 10 days at 37°C. We labeled cells with CD61-PE antibody and collected live cells with the 10% highest and 10% lowest expression of CD61 (**Figure 3A**). sgRNAs in the genomic DNA of each population were sequenced to identify putative genes that affected CD61 expression.

**Figure 3:**
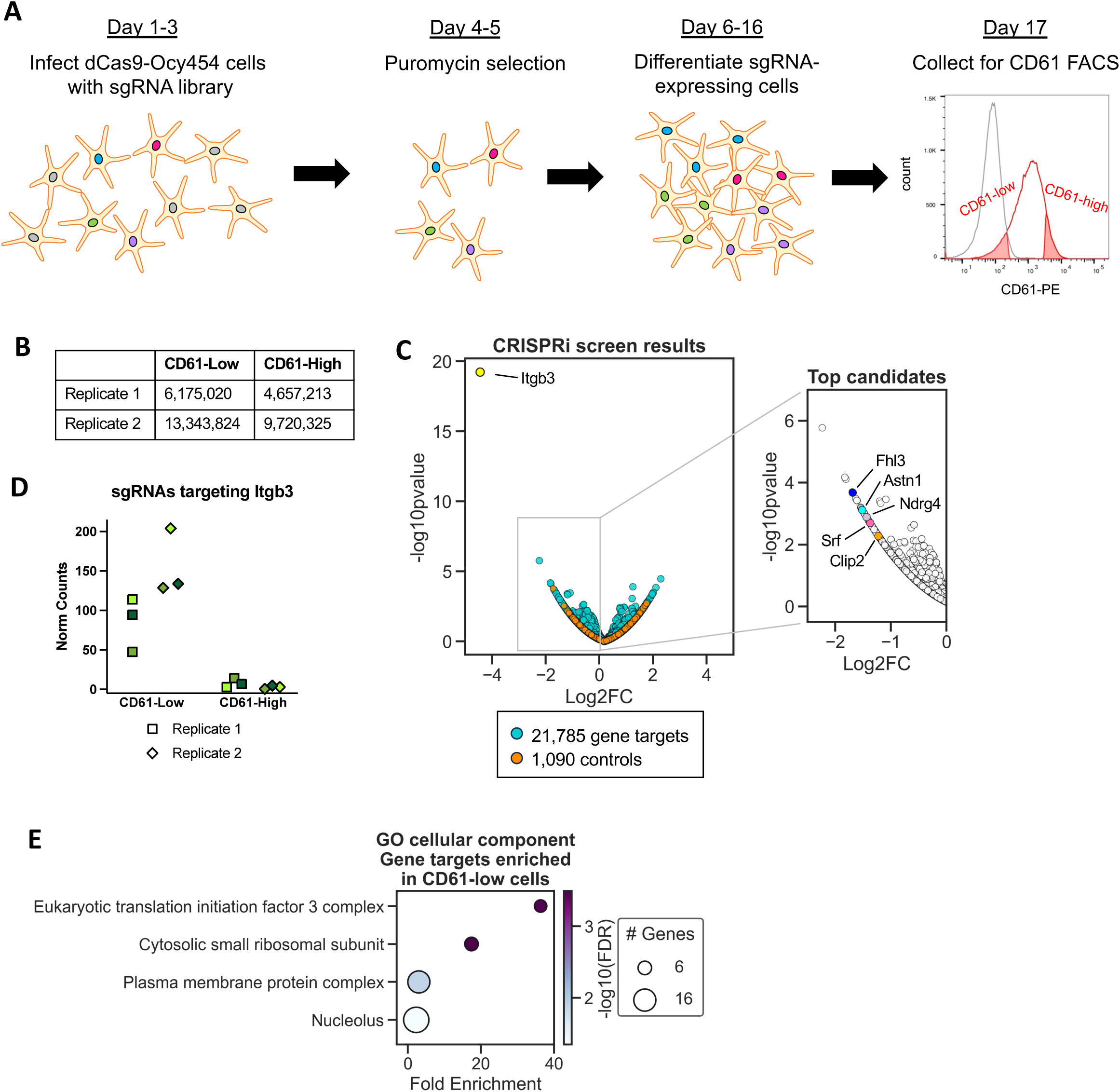
Overview of CRISPRi screening. A) Schematic of CRISPRi screen timeline B) Number of cells collected in CD61^high^ and CD61^low^ groups in each replicate of the CRISPRi screen. C) Volcano plot shows CRISPRi screen results for CD61^high^ vs CD61^low^ groups for both replicates. Each point represents one gene targeted by at least 3 sgRNAs in the pooled screening library. Control points represent groups of three non-targeting or intergenic site-targeting sgRNAs. Inset shows all gene targets enriched in CD61^low^ cells (Log_2_FC<0) with top candidates for follow-up labeled. D) Normalized counts of all sgRNAs targeting the *Itgb3* promoter detected in each replicate of the CRISPRi screen. Colors denote each of three sgRNAs. E) Top gene ontology terms from 202 sgRNAs enriched in CD61-low cells.

In each of two independent replicates, 4.6-13.3 million cells per group were collected for gDNA isolation and sequencing (**Figure 3B**). We compared CD61^low^ to CD61^high^ cells and identified sgRNAs targeting 202 genes significantly enriched in the CD61^low^ group (Log_2_FC<-1, p<0.05) (**Figure 3C, Supp Data 3**). *Itgb3* was the most significantly enriched gene target in the CD61^low^ group in both replicates, which served as a robust positive control (**Figure 3C-D, Supplemental Figure 5D**). Genes encoding components of the translation initiation complex and ribosomal subunit proteins were also enriched in the CD61^low^ group (**Figure 3E, Supplemental Figure 5E-F**). Notably, while complete knockout of genes in these categories would likely have been lethal (https://depmap.org/portal), inhibition with dCas9-KRAB maintains cell viability and demonstrates the necessity of the targeted genes and protein products for cell surface CD61 expression. Additional gene targets enriched in the CD61^low^ group (genes required for CD61 expression) were chosen for further validation, prioritizing non-essential genes with high expression in Ocy454 cells as determined from mRNA-sequencing of this cell line: *Fhl3*, *Astn1*, *Srf*, *Ndrg4*, and *Clip2* (**Figure 3C inset, Supplemental Data 1, Supplemental Data 2**).

### Validation of selected candidates identified in CRISPRi screen via shRNA knockdown

We initially used sgRNAs from the screen library to validate screen results in a non-pooled format, targeting the promoters of *Eif3b*, *Fhl3*, *Astn1*, *Srf*, *Ndrg4*, *Clip2,* and *Itgb3*. However, in doing so we noted decreasing dCas9-KRAB activity with serial passage of dCas9 Ocy454 cells over time. These findings led us to use lentiviral-mediated shRNA knockdown as an orthogonal approach to confirm CRISPRi screening findings (**Supplemental Figure 6A**). Using shRNAs targeting *Eif3b* (**Supplemental Figure 5F**), we observed robust repression of *Eif3b* mRNA, eIF3B protein, and CD61 surface labeling at 3-10 days of differentiation (6-13 days post-shRNA infection) (**Supplemental Figure 6B-D**). Likewise, *Itgb3*-targeting shRNAs reduced both *Itgb3* mRNA and CD61 surface labeling (**Supplemental Figure 6D-G**), validating the use of shRNAs for knockdown in these multi-week experiments.

With *Eif3b* being a common essential gene, we focused on additional gene targets with promoter-targeted sgRNAs enriched in CD61^low^ cells. We designed at least two independent shRNAs for each target and used shRNA sequences targeting firefly luciferase (FLuc), GFP, and LacZ as controls. Nearly all shRNAs reduced mRNA expression of the targeted transcript (**Figure 4A-D**). However, no detectable reduction in *Astn1* expression could be induced with shRNAs, perhaps due to the relatively low baseline expression of this gene (**Supplemental Figure 7A-C**).

**Figure 4.**
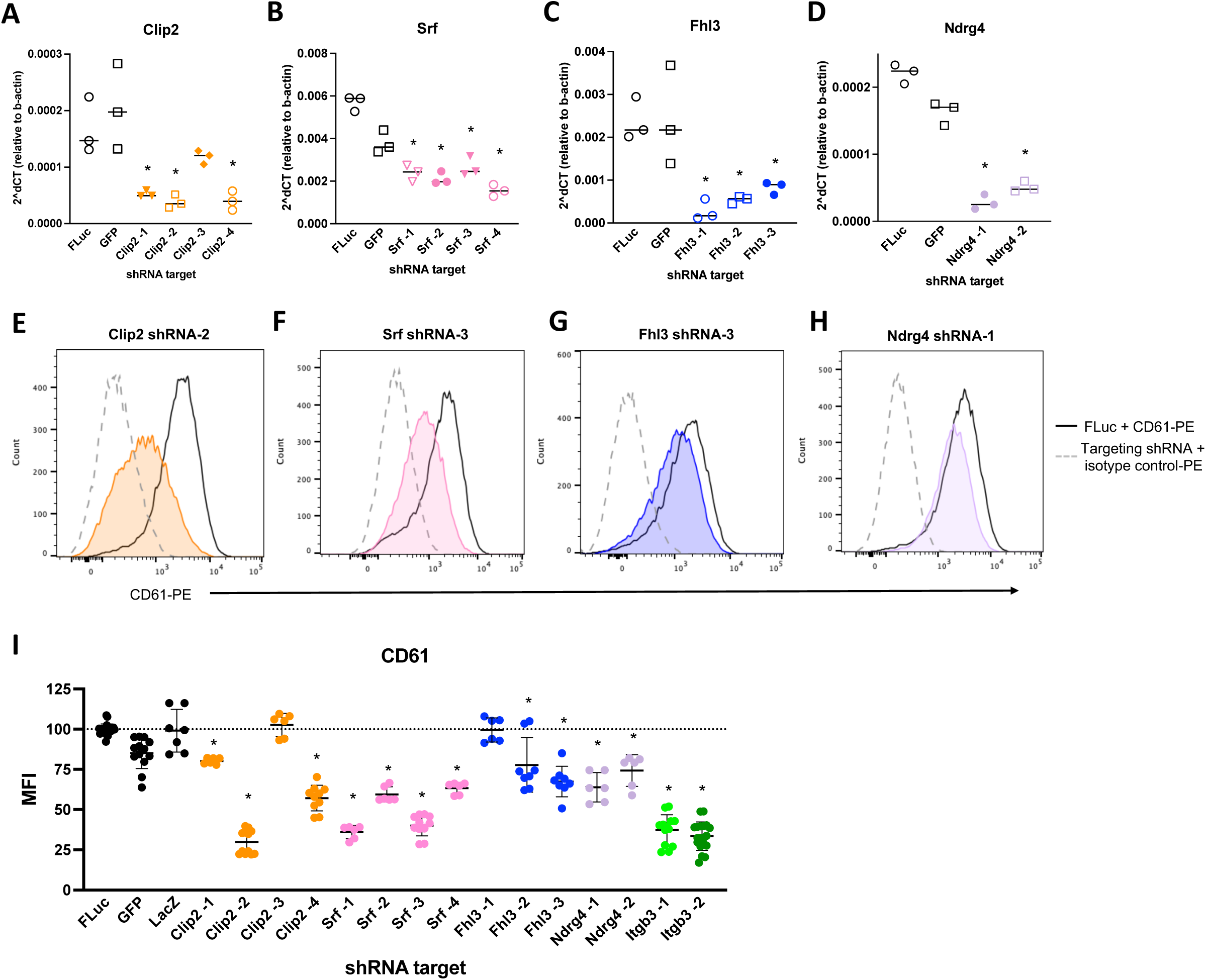
Confirmation of CRISPRi screening hits using shRNA. A-D) mRNA expression of targeted genes at differentiation day 7 in Ocy454 cells stably expressing the indicated shRNAs. Line shows median of three biologic replicates. *p<0.05 compared to FLuc. E-H) Histograms show intensity of CD61-PE fluorescence compared to isotype control at differentiation day 7 for cells stably expressing FLuc-targeting shRNA and the indicated targeting shRNA for one representative sample. I) Summary of flow cytometry results showing median fluorescence intensity (MFI) of CD61-PE at differentiation day 7 in cells stably expressing the indicated shRNAs. Error bars show standard deviation of all replicates. *p<0.05 compared to FLuc.

Across multiple independent experiments, shRNAs targeting *Srf*, *Clip2*, and *Itgb3* reduced CD61 labeling compared to non-targeting control shRNAs. Both the percentage of cells labeled with CD61 (compared to isotype control) and the median fluorescence intensity (normalized to non-targeting shRNAs) were reduced, validating CRISPRi screening results (**Figure 4E-F, I, Supplemental Figure 7D-I**). The magnitude of CD61 inhibition within each gene target was generally related to the extent of knockdown induced by each shRNA. shRNAs targeting *Fhl3* and *Ndrg4* reduced cell MFI but not percentage of cells considered CD61-positive, likely reflecting a more modest overall effect size (**Figure 4G-H, Supplemental Figure 7J-L**).

### Genes that control surface CD61 levels also impact Ocy454 cell maturity and morphology

Next, we assessed the impact of reducing genes that control CD61 levels on other indices of osteocyte maturation. shRNAs targeting *Srf*, *Clip2*, *and Itgb3* significantly reduced expression of osteocyte differentiation markers *Dmp1, Phex,* and *Sost* **(Figure 5A-C, Supplemental Figure 8A-C)**. As with CD61 expression, *Clip2* shRNA knockdown efficiency correlated with the measured phenotype; *Clip2* shRNA-3 was less effective at reducing target mRNA levels and did not reduce osteocyte differentiation gene expression. Also consistent with their more modest effect on CD61, *Ndrg4* and *Fhl3* knockdown did not consistently reduce *Sost* or *Dmp1* expression (**Supplemental Figure 8D-G).**

**Figure 5.**
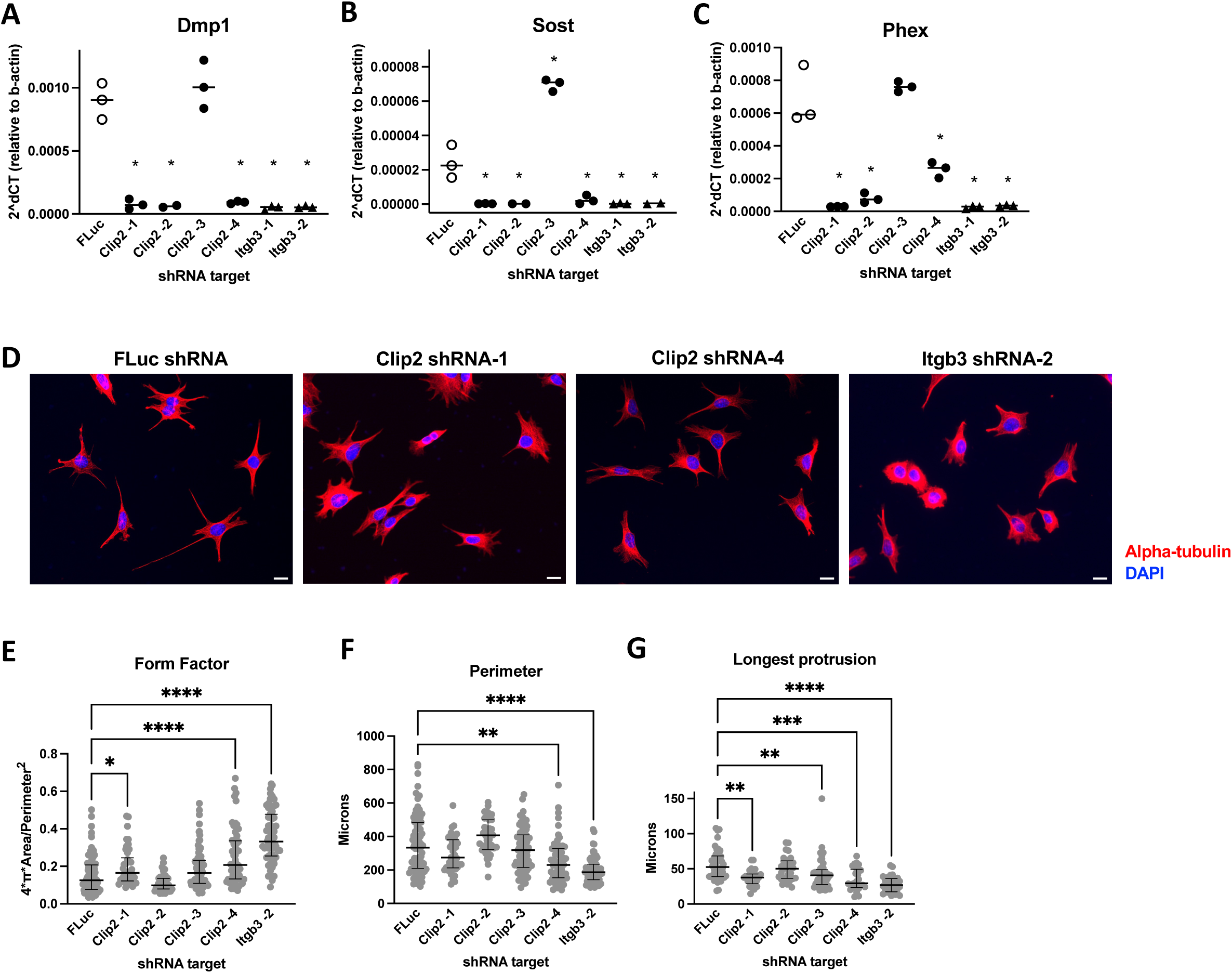
Clip2 knockdown inhibits osteocyte maturation and morphology. A-C) mRNA expression of osteocyte maturity genes at differentiation day 10 in Ocy454 cells stably expressing the indicated shRNAs. Line shows median of biologic replicates. *p<0.05 compared to FLuc. D) Alpha-tubulin immunofluorescence (red) and DAPI (blue) in Ocy454 cells stably expressing the indicated shRNAs. Scale bars = 20 μm. E-F) Cell Profiler measurements of form factor and perimeter. Each plotted point represents metrics of one cell. Lines designate the median and interquartile range of n=36-80 cells. One representative experiment from three independent replicates shown. *p<0.05. F) Quantification of protrusion length for the single longest dendrite of each cell. Lines designate the median and interquartile range of n=24-46 cells. One representative experiment from three independent replicates shown. **p<0.01.

CLIP-115, the protein encoded by *Clip2*, is a microtubule binding protein [39], [40], so we next visualized the microtubule network in cells after knockdown of *Clip2.* Alpha-tubulin immunostaining revealed extensive tubulin structures in the osteocyte cell body and dendrites (**Supplemental Figure 8H**). Ocy454 cells infected with FLuc shRNA have numerous long dendrites with asymmetric distribution (**Figure 5D**). In contrast, osteocyte-like cells expressing *Clip2*-targeting shRNAs have shorter projections, and more round appearance. The overall morphology was quantified as increased (more circular) form factor, reduced perimeter, and reduced maximum projection length, consistent with knockdown of *Itgb3* itself (**Figure 5E-G**). Cell morphology visualized with F-actin staining was also affected by *Clip2* and *Itgb3* knockdown, with significantly increased form factor and reduced perimeter (**Supplemental Figure 8I-K**). Notably, these data suggest that *Itgb3*/CD61 is both a marker and a regulator of osteocyte differentiation.

In summary, a genome-wide CRISPRi screen focused on surface CD61 expression reveals genes with critical regulatory functions in osteocyte differentiation and morphology and demonstrates a role for the microtubule cytoskeleton to drive osteocyte differentiation.

## Discussion

Here we developed an osteocyte maturity assay which allows high-throughput genetic perturbation analysis of the osteocyte differentiation phenotype. CD61 (integrin β3) is identified as a cell surface marker that tracks with differentiation in Ocy454 cells and serves as the basis of a genome-wide CRISPR interference screen. We identify multiple genes required for full surface expression of CD61, including *Clip2*, which is also required for expression of osteocyte differentiation markers and the dendritic osteocyte phenotype. While typically described as actin-based structures, osteocyte dendrites also contain microtubules, which participate in intracellular trafficking, cytoskeletal tension, and osteocyte mechanotransduction [13], [14], [15], [16], [41]. Proper maintenance of these mechanisms may also impact other aspects of gene expression, differentiation, and morphology. Together this work suggests that the microtubule cytoskeleton plays an important role in osteocyte differentiation and morphology.

Many other cell surface markers associated with osteocyte maturity were considered as the readout for our screen. Ideal genome-wide screens require thousands of cells containing each sgRNA to be input into the final sorting of high/low populations [42]. With time required for sorting as a major bottleneck in the workflow, we required a screening readout that labeled the majority of healthy, mature cells and offered a wide dynamic range for detecting sgRNAs causing reduced surface marker intensity. In our experiments, antibodies targeting CXCR7, CA-IX, PTGER4, and CD200 labeled less than 10% of differentiated Ocy454 cells, and CD43 expression lacked the appropriate dynamic range over Ocy454 differentiation time. CD61 is expressed on osteocyte dendrites *in vivo* [32], [33], offering physiological relevance in addition to practicality as an assay readout.

CRISPR screening is an established functional genomics approach to identify gene networks that control distinct cellular functions. Here we chose the dCas9 system in a FACS-based cell differentiation screening readout to avoid potential toxicities of Cas9-induced double strand DNA breaks. Therefore, our screen could capture genes that, when inhibited, interfere directly with *Itgb3* transcription, CD61 translation, or CD61 transport to the cell surface. In addition, since surface CD61 expression is associated with the overall degree of osteocyte maturation, genes that reduce cell surface CD61 levels may also generally participate in other aspects of osteocyte differentiation or general mRNA/protein processing. The latter was apparent in the top Gene Ontology terms enriched in the CD61-low group: gene targets involved generally in ribosome assembly and translation. Since our screen compared two sorted populations of sgRNA-expressing cells, it should not identify genes whose inhibition leads to lethality, which should drop out of all live populations sorted by flow cytometry. We also anticipated that inhibition of most genes would have minimal effect on CD61, and yet identified 202 gene targets significantly enhanced in CD61^low^ cells.

We selected genes for further study based first on enrichment in CD61^low^ cells, and then on their level of mRNA expression in prior Ocy454 cell RNA-seq data and other bone-relevant datasets. Astn1, while one of the most significantly enriched gene targets in the screen, proved challenging to study due to low expression levels and lack of efficient shRNA sequences. Other putative screening ‘hits’ included Fhl3 (binds actin and regulates BMP signaling) [43], [44], Srf (transcription factor that responds to actin dynamics)[45], Ndrg4 (expressed in neurons and in cancer, controls β1-integrin clustering)[46], [47], and Clip2 (microtubule binding protein). Given a previously defined role for Srf in bone mineralization via osteoblast lineage cells [48], we focused our efforts on Clip2.

Clip2, also known as Cyln2 and CLIP-115, is predominantly expressed in the brain, particularly in dendrites within the inferior olive, hippocampus, and piriform cortex [39], [49]. It is one of 28 genes affected in the heterozygous chromosomal deletion that causes Williams Syndrome [50], [51], and its haploinsufficiency likely contributes but is not solely responsible for the neurological symptoms of the disorder [49], [52], [53]. CLIP-115 and closely related protein CLIP-170 (*Clip1*) compete for binding at microtubule plus ends, with CLIP-170 being involved in recruiting the dynein-dynactin complex to microtubule plus ends [49], [54], [55]. Primary fibroblasts isolated from Cyln2 knockout mice showed no measurable differences in microtubule growth rates, however, absence of CLIP-115 resulted in greater localization of CLIP-170 at microtubule plus ends [49], which could contribute to an imbalance in minus-end-directed transport. CLIP-associating proteins (CLASPS) also regulate microtubule dynamics [56], [57]. While *Clip1* inhibition in our screen did not significantly enhance CD61 expression, *Clip2* inhibition is sufficient to disrupt the microtubule network and CD61 surface expression. The role of Clip2 in osteocyte biology *in vivo* represents an important topic for future study.

Early studies of the osteoblast to osteocyte transition visualized microtubules in osteocytes, but concluded that they are only localized in the proximal portion of dendrites and that short-term microtubule depolymerization (60-90 minutes) does not affect osteocyte morphology [11], [12]. These observations contrast with those made for actin; actin and actin-binding proteins are found throughout osteocyte dendrites, and short-term actin depolymerization causes dendrite retraction [11], [12], [58], leading many to focus on the role of actin in these structures [59]. However, we and others observe microtubules throughout osteocyte dendrites *in vitro* and *in situ* [14], [41], and potential roles for microtubules in initiation of dendrite formation and long-term dendrite stability remain largely unexplored. Independent of dendrite macrostructure, microtubules may be required for proper intracellular trafficking [41], [60], [61] and maintenance of mechanosensitive structures [14], [15], [16] to regulate osteocyte function. Furthermore, microtubule modifications and interactions with actin and intermediate filaments can regulate actin polymerization, cytoskeletal tension, and cell motility [62], [63], [64], [65], [66], [67], making additional roles of microtubules and microtubule-binding proteins in osteocytes likely.

In summary, we have used a novel CD61-based Ocy454 cell differentiation assay to identify Clip2 in a genome-wide CRISPR interference screen as an essential contributor to osteocyte morphology and maturation.

## Methods

### Cell culture

Ocy454 osteocyte-like cells [31], [68] were cultured in MEM α supplemented with 10% FBS and 1% antibiotic/antimycotic (Gibco) in a 33°C incubator with 5% CO_2_. At confluence, cells were transferred to a 37°C incubator to induce differentiation.

Ocy454 cells were plated at 40,000 cells/mL media the day before lentiviral infection. Fresh media containing 8 µg/mL polybrene and lentiviral particles was added the next day and allowed to incubate overnight. Media was changed the next day to remove the transduction reagents, and then again the next day to begin puromycin selection (4 µg/mL). Cells receiving no lentivirus with or without puromycin were used as infection controls. When puromycin-resistant cells reached confluence they were transferred to a 37°C incubator (differentiation day 0).

Sp7 deficient Ocy454 cells stably express a Sp7-targeting shRNA [5]. HDAC4/HDAC5 double knockout Ocy454 cells stably express a HDAC4-targeting sgRNA and an HDAC5-targeting shRNA [36].

### dCas9-KRAB osteocytes

Ocy454 cells were infected with lentivirus containing UCOE-SFFV-dCas9-BFP-KRAB (Addgene #85969) [38] and expanded in culture for 3 days. BFP positive cells were selected by flow cytometry, cultured for another 3 days, and re-sorted for BFP. The resulting polyclonal cell population was used for all sgRNA experiments.

### shRNA and sgRNA preparation

CRISPRi sgRNA sequences and shRNA sequences were designed using Broad Institute tools CRISPick (https://portals.broadinstitute.org/gppx/crispick/public) and TRC shRNA Design (https://portals.broadinstitute.org/gpp/public/seq/search). All sgRNA and shRNA sequences are available in Supplemental Data 4. sgRNAs were ligated into pLentiGuide (Addgene #117986) and shRNAs were ligated into pLKO.1 (Addgene #10878) using standard protocols, and clones validated by Sanger sequencing.

### Lentivirus preparation

HEK293T cells were cultured in DMEM supplemented with 10% FBS and 0.1% antibiotic/antimycotic in at 37°C incubator with 5% CO_2_. Cells were plated into 6-well tissue culture plates at 440,000 cells/well. The next day, cells were transfected with viral packaging plasmids psPAX2 and pMD2.G (Addgene #12260, #12259) and insert plasmids containing a single sgRNA or shRNA using PolyJet reagent (SignaGen Laboratories SL100688). Lentivirus was collected for 48 hours in media containing 20% FBS, then stored in aliquots at -80°C.

### mRNA isolation

Ocy454 cells were plated into 24-well plates, individually infected with lentivirus, and differentiated at 37°C for 7 days. Cells were then washed in PBS and lysed in RLT lysis buffer. Lysates were passed through QIAshredder columns (Qiagen), then purified with PureLink RNA columns (Invitrogen) according to manufacturer’s instructions.

### qRT-PCR

Total RNA was reverse transcribed to cDNA with PrimeScript RT Reagent Kit (Takara, including gDNA digestion step). qPCR was performed using SYBR green with CT values normalized to *b-actin* (Quantabio). All primer sequences are in Supplemental Data 4.

### Flow cytometry

Ocy454 cells at the specified differentiation timepoint were collected with 0.5% Trypsin-EDTA, washed in FACS buffer (2% FBS in PBS), and resuspended in FACS buffer at approximately 1M cells/mL. Cells were incubated in FACS buffer at 4°C for 30 minutes. Individual reactions were prepared containing approximately 100,000 cells each with 0.5 µL of anti-CD61 (Invitrogen 12-0611-82) or isotype control (Invitrogen 12-4888-83) antibody and incubated at 4°C for 45 minutes with gentle rocking. Cells were then washed in FACS buffer, resuspended in FACS buffer, and labeled with SYTOX green dead cell stain before flow cytometry. Flow cytometry analysis was primarily performed on an Attune NxT flow cytometer.

### Differentiation RNA-seq

Ocy454 cells were differentiated for 1, 7, 10, or 14 days at 37°C, then collected for RNA in biologic duplicates, as described. Total RNA was used for library preparation and sequencing (BGI DNBseq platform). STAR aligner was used to map sequencing reads to transcripts in the mouse reference genome [69]. Read counts for individual transcripts were produced with HTSeq-count [70], followed by estimation of expression values using EdgeR. Differential expression analysis was performed using EdgeR after normalizing read counts and including only genes with count per million (CPM) reads >1 for at least one sample.

### CD61 RNA-seq

Ocy454 cells were differentiated for 17 days at 37°C. Three independent groups of cells were then collected and prepared for CD61 flow cytometry as described. The top 10% and bottom 10% of live CD61-PE labeled cells were collected into FACS buffer containing RNase inhibitor. Cells were then pelleted and lysed in RLT buffer. Lysates were passed through QIAshredder columns, then total RNA was isolated using the Arcturus PicoPure RNA isolation kit (Applied Biosystems) with on-column DNase (QIAgen). Total RNA was used for library preparation and sequencing (BGI DNBseq platform).

At least 18 million paired-end reads per sample were obtained from PolyA-selected libraries, with >89% of reads mapping uniquely to the mouse transcriptome (GRCm39 Release 104; Salmon)[71]. Transcript counts were collapsed to gene counts, and differential expression analysis was performed with DESeq2 [72]. Genes with at least 5 normalized counts in all samples were considered for analysis. RNA sequencing data will be deposited at NCBI’s Gene Expression Omnibus and made available upon request to the corresponding author.

### CRISPRi screen

The Dolomiti A lentivirus library [73] containing 67,366 sgRNAs targeting promoters of mouse genes and control sequences (approximately 3 sgRNAs/gene target) was purchased from the Broad Institute. Viral titer was initially determined by measuring puromycin resistance following serial dilutions of virus in Ocy454 cells. Two independent replicates of the screen were performed. 24 million cells in 15 15-cm tissue culture dishes were infected at target MOI 0.3-0.5 in MEM-alpha media containing 8 µg/mL polybrene (Sigma H9268). On day 2 post-infection, sgRNA-containing cells were selected with puromycin (4 µg/mL). At confluence, puromycin-resistant cells were transferred to 37°C to induce differentiation. On differentiation day 10, cells were collected in three batches and labeled with CD61-PE and SYTOX-Green as described above. Sorting was performed on a FACSAria (Becton Dickinson) to collect the top 10% and bottom 10% of the live CD61-PE-labeled population. Sorted cells were collected into 100% FBS prior to genomic DNA isolation.

### CRISPRi screen analysis

Cell pellets from each group were stored at -80°C, then genomic DNA (gDNA) was extracted with the NucleoSpin Blood Kit (Machery Nagel) with the following modifications from the manufacturer’s instructions: lyse cell pellets overnight, then treat with RNase A for 5 minutes. After DNA precipitation and washing, further purify the gDNA with PCR inhibitor removal kit (Zymo #D6030). sgRNA sequences were PCR amplified from gDNA using Titanium *Taq* DNA Polymerase. 4 replicate reactions per sample each containing 10 µg gDNA were amplified with 32 PCR cycles. Primers added barcodes for each reaction and Illumina sequencing adaptors. PCR products were purified, pooled, and sequenced in one lane of a HiSeq-2500 (Illumina).

Reaction barcodes and sgRNAs were parsed from each sequencing read using PoolQ, then counts were assigned to known library sgRNAs without allowing mismatches. Total sgRNA counts for all reactions for each sample were then aggregated. Target-level aggregation, fold change calculations, and statistical analysis for both replicates were performed with Apron for all gene targets with at least three sgRNAs. Non-targeting and intergenic site-targeting sgRNAs were grouped into 3-guide pseudogenes for points plotted as controls.

### Western blots

Cells expressing shRNAs were washed in cold PBS, pelleted, and lysed using TNT protein lysis buffer (20 mM Tris pH 8, 200 mM NaCl, 0.5% Triton X-100) supplemented with NaF, vanadate, DTT, and protease inhibitor cocktail (Selleck USA). Clarified proteins were separated on 8% SDS-PAGE gels, blocked, and labeled overnight with antibodies against eIF3B (Bethyl A301-760A) and Gapdh (Cell Signaling #2118), then detected with HRP-linked anti-rabbit secondary developed with ECL Plus (Thermo Scientific Pierce).

### Immunofluorescence

Ocy454 cells expressing shRNAs were plated into chamber slides and allowed to propagate for at least two days. Cells were then fixed in pre-warmed 4% PFA for 15 minutes, washed in PBS, and permeabilized in 0.5% Triton X-100 in HBSS. F-actin staining (Abcam Cytopainter Green) was performed at room temperature for 1-2 hours, followed by DAPI labeling.

For alpha tubulin immunofluorescence, permeabilized cells were blocked in 5% BSA in HBSS with 0.1% Triton X-100, then incubated with anti-alpha tubulin (Sigma T9026) or mouse IgG (sc-2025, 2.6 µg/mL final concentration) overnight at 4°C [14], [74]. Secondary antibody (Invitrogen A31570) was applied in 1% BSA in HBSS with 0.1% Triton X-100 or with F-actin staining buffer at room temperature for 2 hours, followed by DAPI labeling. Imaging was performed on a Nikon Eclipse microscope.

### Image analysis

Single-channel images were analyzed with CellProfiler (v4.2.6) to identify single cells and measure quantitative morphology features [75]. Our pipeline segmented DAPI-stained nuclei as primary objects, then used the phalloidin-stained actin cytoskeleton or the alpha tubulin-stained microtubule cytoskeleton to segment individual cells. Form factor and perimeter were automatically measured for each identified cell. Protrusion length from the edge of the nucleus to the tip was measured manually for the longest dendrite of each cell in ImageJ [76].

### Statistical analysis

With the exception of sequencing-based outcomes, statistical analyses were performed in GraphPad Prism v10.0.2. Intensity and proportion of CD61-labeling was compared by one-way ANOVA with Dunnett’s multiple comparisons tests against no sgRNA control or against FLuc-targeting shRNA. qPCR expression of target genes was compared by one-way ANOVA with Dunnett’s multiple comparisons tests against cells infected with FLuc-targeting shRNA. Cell morphology metrics were analyzed with one-way ANOVA on ranks followed by Dunn’s multiple comparisons tests.

## Acknowledgements

We thank all members of the MGH Endocrine Unit and the Wein laboratory for helpful discussions. We thank Ruslan Sandreyev and the Bioinformatics Core of the MGH Department of Molecular Biology for assistance with analysis of the Ocy454 differentiation RNA-sequencing. M.N.W. acknowledges funding support from the National Institute of Health (R01DK116716), the Smith Family Foundation Odyssey Award, and the Chen Institute Massachusetts General Hospital Research Scholar (2024-2029) award. T.S acknowledges support from NIH/NIAMS (R21AR079633, R21AR084644) and Center for Skeletal Research P30 core P&F award (P30 AR075042). C.M.M. acknowledges support from the NIH (T32DK007028, F32AR081660). M.N.W. and C.M.M. acknowledge generous support from Louise Pearl Corman, Ph.D. Fluorescence-activated cell sorting was performed at the Wellman Center for Photomedicine core.

## Author contributions

Conceptualization: C.M.M. and M.N.W.

Methodology: J.D.

Investigation: C.M.M., P.E.K., M.G., T.W.

Resources: L.E.S., T.S.

Writing: C.M.M. and M.N.W.

Funding Acquisition: M.N.W.

## Supplementary Information

Document S1: Supplemental Figures 1-8.

Supplemental Data 1: Table of normalized gene counts and differential expression from Ocy454 differentiation RNA sequencing.

Supplemental Data 2: Table of normalized gene counts and differential expression from CD61 FACS RNA sequencing.

Supplemental Data 3: List of gene targets significantly enriched (Log_2_FC<-1, p<0.05) in the CD61^low^ group of CRISPRi screen.

Supplemental Data 4: Primers and DNA oligos.

---

**Supplemental Figure 1:**
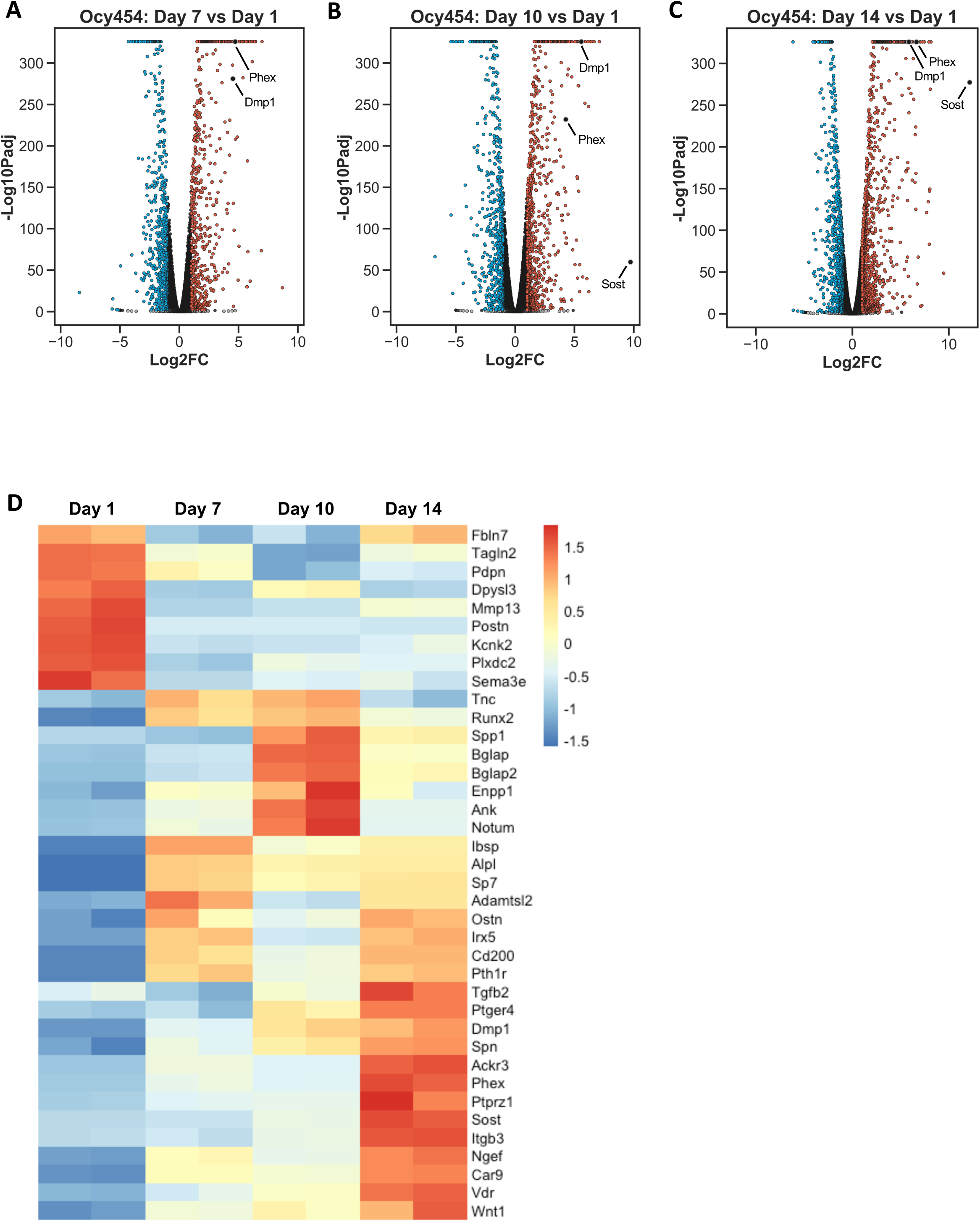
A) Volcano plot of Ocy454 differentiation at day 7 compared to day 1. B) Volcano plot of Ocy454 differentiation at day 10 compared to day 1. C) Volcano plot of Ocy454 differentiation at day 14 compared to day 1. D) Heat map shows z-scored counts per million for osteoblast to osteocyte transition genes, osteocyte signature genes, and genes encoding prospective surface markers for flow cytometry. All genes included in heat map are significantly differentially expressed (FDR<0.01) in at least one comparison to Day 1. Scaling is calculated individually for each row; therefore, no inferences should be made about relative abundances of genes at each timepoint.

**Supplemental Figure 2:**
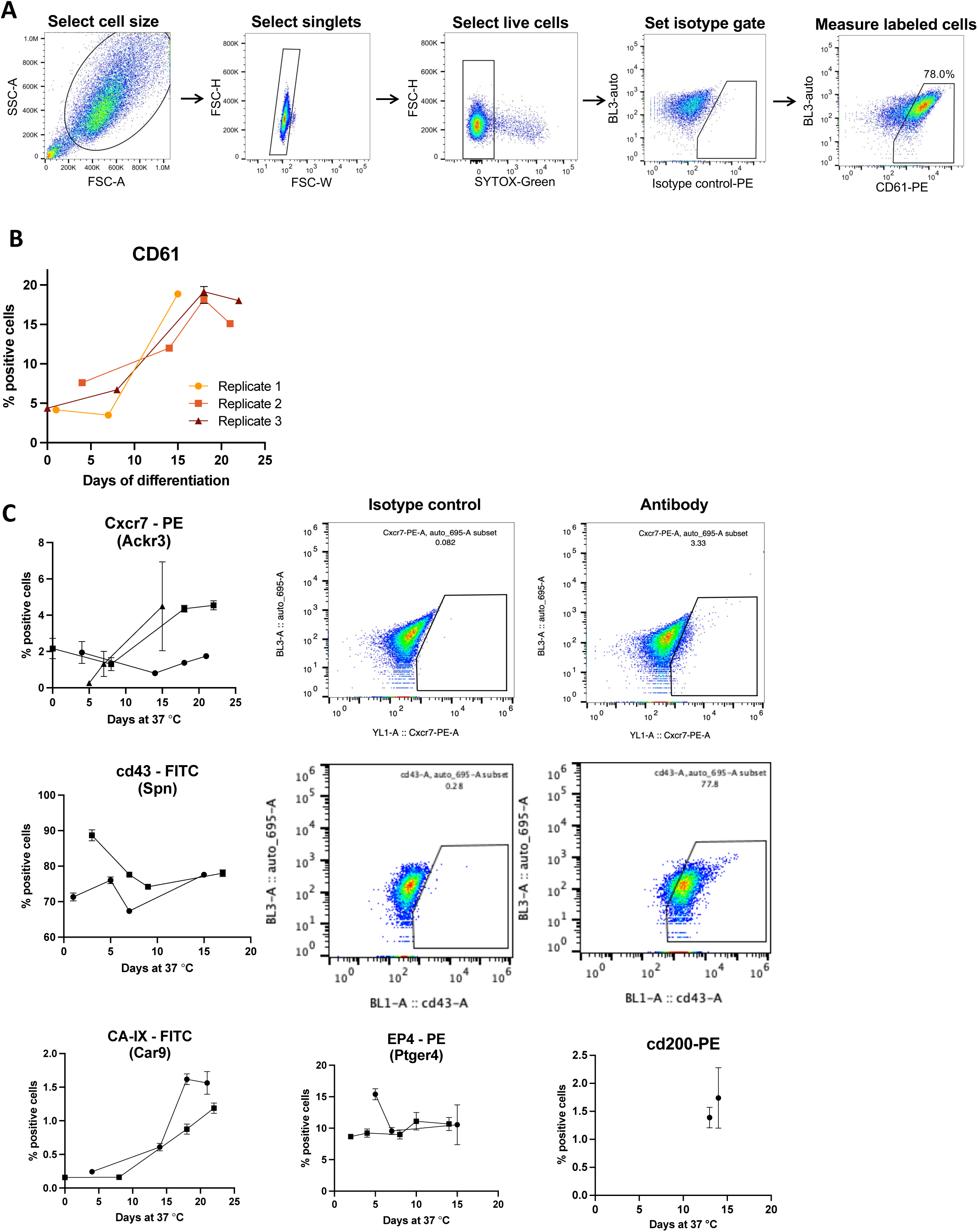
A) Flow cytometry gating strategy for CD61 detection in live Ocy454 cells B) Flow cytometry results show percent of cells labeled with CD61-FITC antibody compared to isotype control at a range of differentiation timepoints. Lines connect matched subcultures of cells followed over time. Error bars show standard deviation for technical replicates in each test. C) Flow cytometry results show percent of Ocy454 cells labeled with indicated antibodies compared to matched isotype controls at a range of differentiation timepoints. Lines connect matched subcultures of cells followed over time. Error bars show standard deviation for technical replicates in each assay.

**Supplemental Figure 3:**
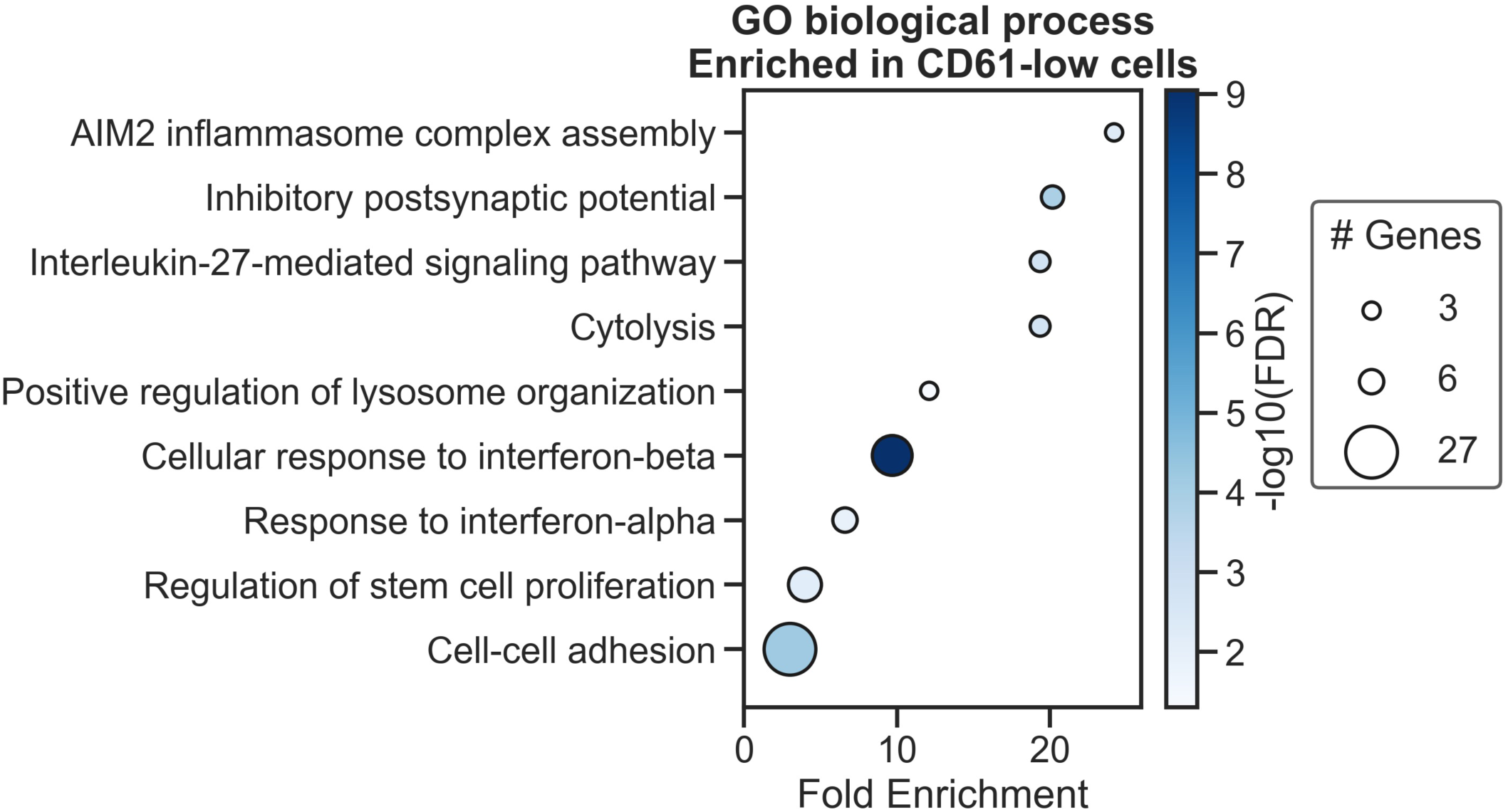
Top gene ontology terms of 501 genes enriched in CD61-low cells (Log_2_FC>1 and p_adj_<0.01).

**Supplemental Figure 4:**
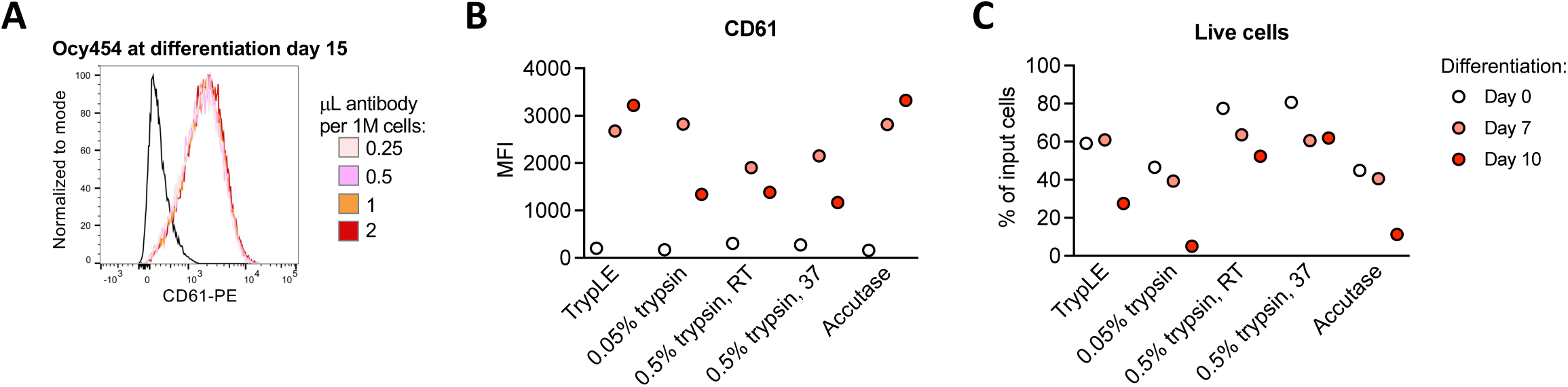
A) Histograms show intensity of CD61-PE fluorescence compared to isotype control (black line) when equal numbers of cells were incubated with a range of total antibody. B) Median fluorescence intensity (MFI) of CD61-PE following cell dissociation with each reagent at a range of differentiation timepoints. Each point represents one test of >25,000 live single cells. C) Percent of prepared cells passing FACS gates based on size, singlets, and viability following cell dissociation with each reagent at a range of differentiation timepoints.

**Supplemental Figure 5:**
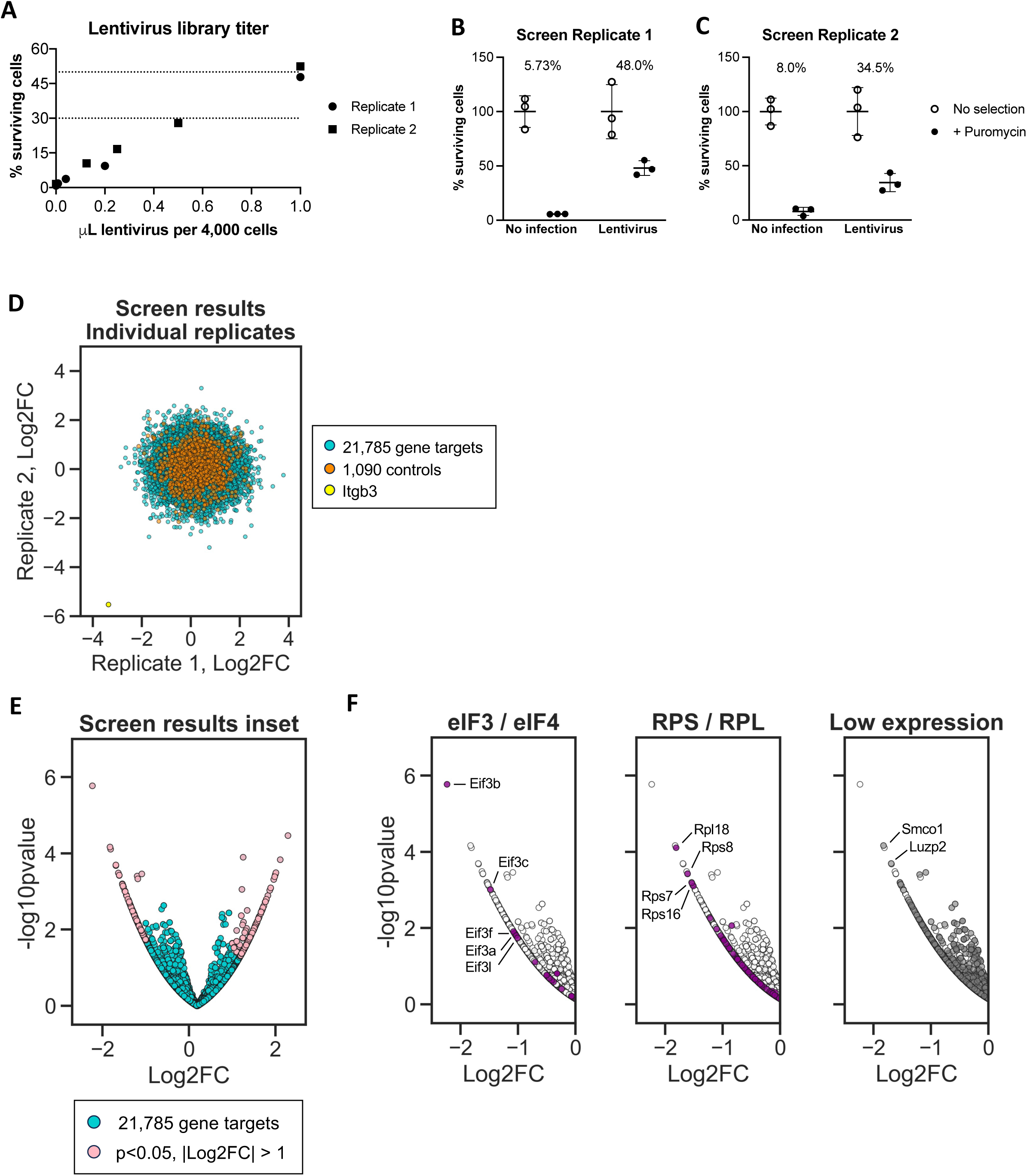
A) Scatter plot showing cell viability after infection with Dolomiti A lentivirus library and puromycin selection, indicating multiplicity of infection (MOI) achieved with each volume of lentivirus. The targeted MOI of 30-50% is indicated with horizontal dashed lines. B) Cell viability after infection with Dolomiti A lentivirus library or no lentivirus and puromycin selection or no puromycin selection in CRISPRi screen replicate 1. Cells were treated in parallel to the screen and prepared in biologic triplicates. Counts were performed after 5 days of puromycin selection. C) Cell viability after infection with Dolomiti A lentivirus library or no lentivirus and puromycin selection or no puromycin selection in CRISPRi screen replicate 2. Cells were treated in parallel to the screen and prepared in biologic triplicates. Counts were performed after 4 days of puromycin selection. D) Scatter plot shows Log2FC results for CD61^high^ vs CD61^low^ in two individual replicates of the CRISPRi screen. Each point represents one gene targeted by at least 3 sgRNAs in the pooled screening library. Control points represent groups of three non-targeting or intergenic site-targeting sgRNAs. E) Inset of volcano plot showing CRISPRi screen results for CD61^high^ vs CD61^low^ groups for both replicates. Control sgRNAs and Itgb3-targeting sgRNAs are omitted. Each point represents one gene targeted by at least 3 sgRNAs in the pooled screening library. F) Volcano plot insets showing all gene targets enriched in CD61^low^ cells (Log_2_FC<0). Labeled genes encode components of the eukaryotic translation initiation factor complex (eIF3/eIF4), encode small or large ribosomal protein subunits (RPS/RPL), or are not detected in RNA-sequencing datasets of Ocy454 cells (Supplemental Figure 1, Figure 1G).

**Supplemental Figure 6:**
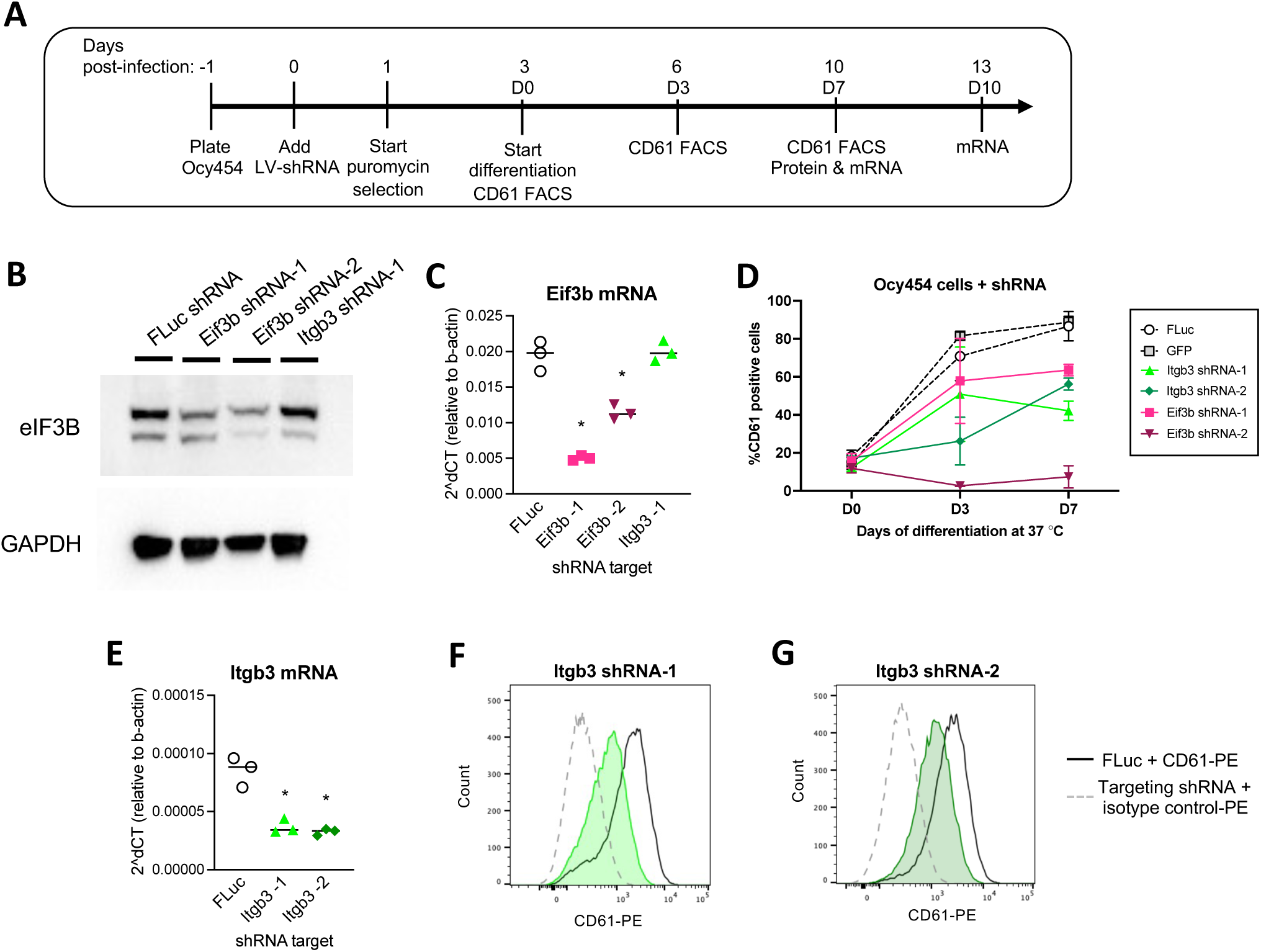
A) Timeline of shRNA infection and analysis of mRNA and protein expression. B) Western blot shows expression of eIF3B protein in Ocy454 cells at differentiation day 7 following stable infection with the indicated shRNAs. C) mRNA expression of *Eif3b* in Ocy454 cells at differentiation day 7 following stable infection with the indicated shRNAs. Line shows median of three biologic replicates. *p<0.05. D) Flow cytometry results show percent of cells labeled with CD61-PE antibody compared to isotype control at a range of differentiation timepoints following stable infection with the indicated shRNAs. Error bars show standard deviation for all replicates at each time point. E) mRNA expression of *Itgb3* in Ocy454 cells at differentiation day 10 following stable infection with the indicated shRNAs. Line shows median of three biologic replicates. *p<0.05. F) Histograms show intensity of CD61-PE fluorescence compared to isotype control at differentiation day 7 for cells stably expressing FLuc-targeting shRNA and Itgb3-targeting shRNA. G) Histograms show intensity of CD61-PE fluorescence compared to isotype control at differentiation day 7 for cells stably expressing FLuc-targeting shRNA and Itgb3-targeting shRNA.

**Supplemental Figure 7:**
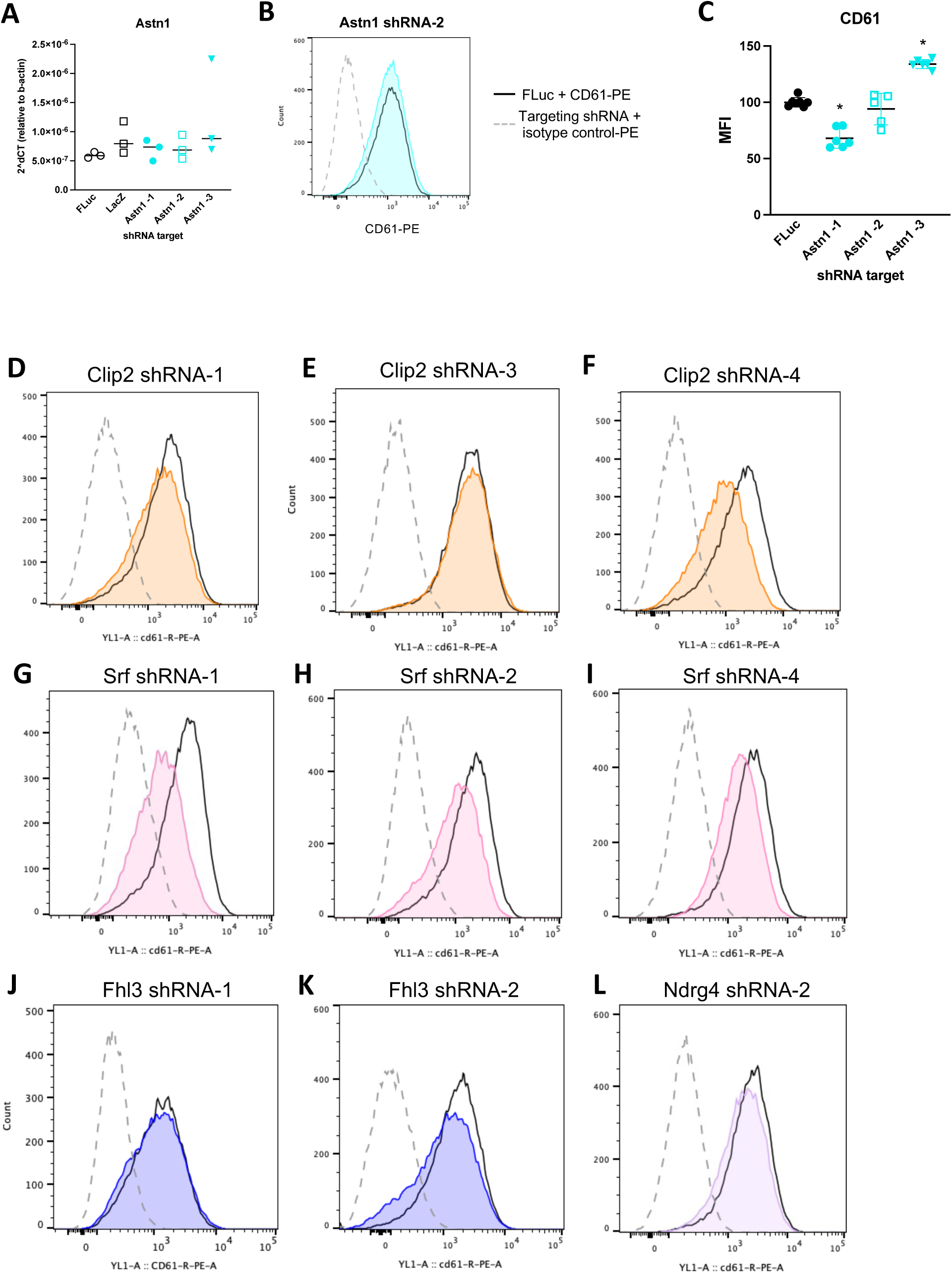
A) mRNA expression of *Astn1* in Ocy454 cells at differentiation day 7 following stable infection with the indicated shRNAs. Line shows median of three biologic replicates. B) Histograms show intensity of CD61-PE fluorescence compared to isotype control at differentiation day 7 for cells stably expressing FLuc-targeting shRNA and Astn1-targeting shRNA for one representative sample. C) Summary of flow cytometry results showing median fluorescence intensity (MFI) of CD61-PE at differentiation day 7 in cells stably expressing the indicated shRNAs. Error bars show standard deviation of all replicates. *p<0.05 compared to FLuc. D-L) Histograms show intensity of CD61-PE fluorescence compared to isotype control at differentiation day 7 for cells stably expressing FLuc-targeting shRNA and the indicated targeting shRNA for one representative sample.

**Supplemental Figure 8:**
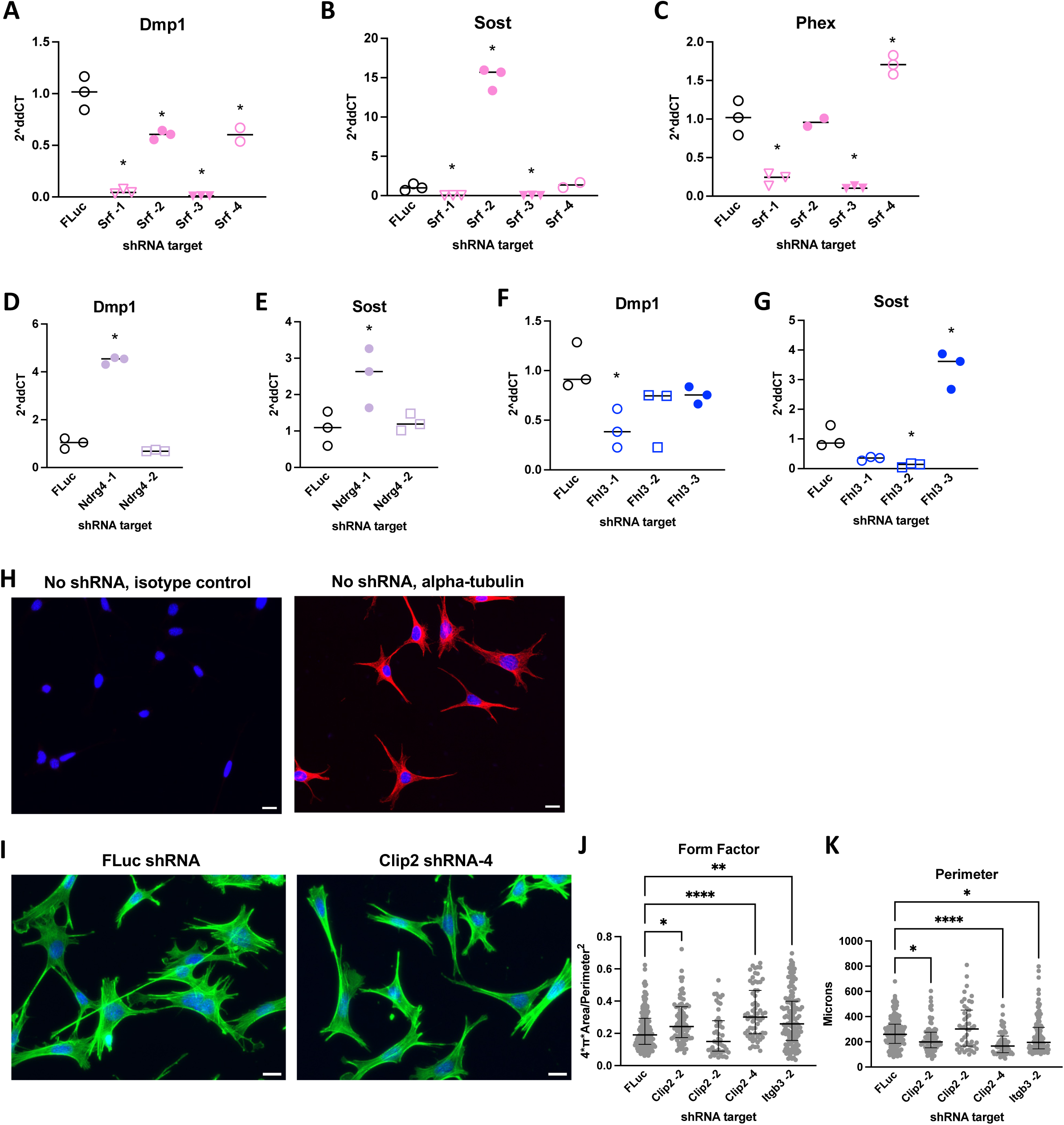
A-G) mRNA expression of osteocyte maturity genes at differentiation day 10 in Ocy454 cells stably expressing the indicated shRNAs. *p<0.05 compared to FLuc. H) Alpha-tubulin immunofluorescence or isotype control immunofluorescence (red) and DAPI (blue) in Ocy454 cells without shRNA infection. Scale bars = 20 μm. I) Ocy454 cells labeled with phalloidin (green) and DAPI (blue) following stable expression of the indicated shRNAs. Scale bars = 20 μm. J-K) Cell Profiler measurements of form factor and perimeter. Each plotted point represents metrics of one cell. Lines designate the median and interquartile range of n=47-158 cells. *p<0.05.

## References

[1] S. L. Dallas, M. Prideaux, and L. F. Bonewald, “The osteocyte: an endocrine cell … and more,” Endocr Rev, vol. 34, no. 5, pp. 658–690, Oct. 2013, doi: 10.1210/er.2012-1026.

[2] M. B. Schaffler, W.-Y. Cheung, R. Majeska, and O. Kennedy, “Osteocytes: master orchestrators of bone,” Calcif Tissue Int, vol. 94, no. 1, pp. 5–24, Jan. 2014, doi: 10.1007/s00223-013-9790-y.

[3] S. Tatsumi et al., “Targeted ablation of osteocytes induces osteoporosis with defective mechanotransduction,” Cell Metab, vol. 5, no. 6, pp. 464–475, June 2007, doi: 10.1016/j.cmet.2007.05.001.

[4] A. G. Robling and L. F. Bonewald, “The Osteocyte: New Insights,” Annu Rev Physiol, vol. 82, pp. 485–506, Feb. 2020, doi: 10.1146/annurev-physiol-021119-034332.

[5] J. S. Wang et al., “Control of osteocyte dendrite formation by Sp7 and its target gene osteocrin,” Nat Commun, vol. 12, no. 1, p. 6271, Nov. 2021, doi: 10.1038/s41467-021-26571-7.

[6] S. E. Youlten et al., “Osteocyte transcriptome mapping identifies a molecular landscape controlling skeletal homeostasis and susceptibility to skeletal disease,” Nat Commun, vol. 12, no. 1, p. 2444, May 2021, doi: 10.1038/s41467-021-22517-1.

[7] B. Cheng, S. Zhao, J. Luo, E. Sprague, L. F. Bonewald, and J. X. Jiang, “Expression of functional gap junctions and regulation by fluid flow in osteocyte-like MLO-Y4 cells,” J Bone Miner Res, vol. 16, no. 2, pp. 249–259, Feb. 2001, doi: 10.1359/jbmr.2001.16.2.249.

[8] S. B. Doty, “Morphological evidence of gap junctions between bone cells,” Calcif Tissue Int, vol. 33, no. 5, pp. 509–512, 1981, doi: 10.1007/BF02409482.

[9] P. R. Buenzli and N. A. Sims, “Quantifying the osteocyte network in the human skeleton,” Bone, vol. 75, pp. 144–150, June 2015, doi: 10.1016/j.bone.2015.02.016.

[10] L. Qin, W. Liu, H. Cao, and G. Xiao, “Molecular mechanosensors in osteocytes,” Bone Res, vol. 8, p. 23, 2020, doi: 10.1038/s41413-020-0099-y.

[11] S. A. Murshid, H. Kamioka, Y. Ishihara, R. Ando, Y. Sugawara, and T. Takano-Yamamoto, “Actin and microtubule cytoskeletons of the processes of 3D-cultured MC3T3-E1 cells and osteocytes,” J Bone Miner Metab, vol. 25, no. 3, pp. 151–158, 2007, doi: 10.1007/s00774-006-0745-5.

[12] K. Tanaka-Kamioka, H. Kamioka, H. Ris, and S.-S. Lim, “Osteocyte Shape Is Dependent on Actin Filaments and Osteocyte Processes Are Unique Actin-Rich Projections,” Journal of Bone and Mineral Research, vol. 13, no. 10, pp. 1555–1568, Oct. 1998, doi: 10.1359/jbmr.1998.13.10.1555.

[13] N. R. Gould, O. M. Torre, J. M. Leser, and J. P. Stains, “The cytoskeleton and connected elements in bone cell mechano-transduction,” Bone, vol. 149, p. 115971, Aug. 2021, doi: 10.1016/j.bone.2021.115971.

[14] J. S. Lyons et al., “Microtubules tune mechanotransduction through NOX2 and TRPV4 to decrease sclerostin abundance in osteocytes,” Sci Signal, vol. 10, no. 506, p. eaan5748, Nov. 2017, doi: 10.1126/scisignal.aan5748.

[15] J. G. McGarry, J. Klein-Nulend, and P. J. Prendergast, “The effect of cytoskeletal disruption on pulsatile fluid flow-induced nitric oxide and prostaglandin E2 release in osteocytes and osteoblasts,” Biochem Biophys Res Commun, vol. 330, no. 1, pp. 341–348, Apr. 2005, doi: 10.1016/j.bbrc.2005.02.175.

[16] L. I. Plotkin, I. Mathov, J. I. Aguirre, A. M. Parfio, S. C. Manolagas, and T. Bellido, “Mechanical stimulation prevents osteocyte apoptosis: requirement of integrins, Src kinases, and ERKs,” Am J Physiol Cell Physiol, vol. 289, no. 3, pp. C633–643, Sept. 2005, doi: 10.1152/ajpcell.00278.2004.

[17] H. C. St. John et al., “The Osteoblast to Osteocyte Transition: Epigenetic Changes and Response to the Vitamin D_3_ Hormone,” Molecular Endocrinology, vol. 28, no. 7, pp. 1150–1165, July 2014, doi: 10.1210/me.2014-1091.

[18] O. Shalem et al., “Genome-Scale CRISPR-Cas9 Knockout Screening in Human Cells,” Science, vol. 343, no. 6166, pp. 84–87, Jan. 2014, doi: 10.1126/science.1247005.

[19] O. O. Abudayyeh et al., “RNA targeting with CRISPR-Cas13,” Nature, vol. 550, no. 7675, pp. 280–284, Oct. 2017, doi: 10.1038/nature24049.

[20] L. Cong et al., “Multiplex Genome Engineering Using CRISPR/Cas Systems,” Science, vol. 339, no. 6121, pp. 819–823, Feb. 2013, doi: 10.1126/science.1231143.

[21] L. A. Gilbert et al., “CRISPR-mediated modular RNA-guided regulation of transcription in eukaryotes,” Cell, vol. 154, no. 2, pp. 442–451, July 2013, doi: 10.1016/j.cell.2013.06.044.

[22] M. Han et al., “Programmable control of spatial transcriptome in live cells and neurons,” Nature, vol. 643, no. 8070, pp. 241–251, July 2025, doi: 10.1038/s41586-025-09020-z.

[23] A. C. Komor, Y. B. Kim, M. S. Packer, J. A. Zuris, and D. R. Liu, “Programmable editing of a target base in genomic DNA without double-stranded DNA cleavage,” Nature, vol. 533, no. 7603, pp. 420–424, May 2016, doi: 10.1038/nature17946.

[24] J. M. Baronas et al., “Genome-wide CRISPR screening of chondrocyte maturation newly implicates genes in skeletal growth and height-associated GWAS loci,” Cell Genom, vol. 3, no. 5, p. 100299, May 2023, doi: 10.1016/j.xgen.2023.100299.

[25] L. Bassaganyas et al., “New factors for protein transport identified by a genome-wide CRISPRi screen in mammalian cells,” J Cell Biol, vol. 218, no. 11, pp. 3861–3879, Nov. 2019, doi: 10.1083/jcb.201902028.

[26] J. M. Dempster et al., “Agreement between two large pan-cancer CRISPR-Cas9 gene dependency data sets,” Nat Commun, vol. 10, no. 1, p. 5817, Dec. 2019, doi: 10.1038/s41467-019-13805-y.

[27] K. M. C. Klee et al., “A CRISPR screen in intestinal epithelial cells identifies novel factors for polarity and apical transport,” Elife, vol. 12, p. e80135, Jan. 2023, doi: 10.7554/eLife.80135.

[28] M. Ramezani et al., “A genome-wide atlas of human cell morphology,” Nat Methods, vol. 22, no. 3, pp. 621–633, Mar. 2025, doi: 10.1038/s41592-024-02537-7.

[29] C. H. Wong et al., “Genome-scale requirements for dynein-based transport revealed by a high-content arrayed CRISPR screen,” J Cell Biol, vol. 223, no. 5, p. e202306048, May 2024, doi: 10.1083/jcb.202306048.

[30] K. You et al., “QRICH1 dictates the outcome of ER stress through transcriptional control of proteostasis,” Science, vol. 371, no. 6524, p. eabb6896, Jan. 2021, doi: 10.1126/science.abb6896.

[31] M. N. Wein et al., “HDAC5 controls MEF2C-driven sclerostin expression in osteocytes,” J Bone Miner Res, vol. 30, no. 3, pp. 400–411, Mar. 2015, doi: 10.1002/jbmr.2381.

[32] P. Cabahug-Zuckerman et al., “Potential role for a specialized β3 integrin-based structure on osteocyte processes in bone mechanosensation,” J Orthop Res, vol. 36, no. 2, pp. 642–652, Feb. 2018, doi: 10.1002/jor.23792.

[33] L. M. McNamara, R. J. Majeska, S. Weinbaum, V. Friedrich, and M. B. Schaffler, “Aoachment of osteocyte cell processes to the bone matrix,” Anat Rec (Hoboken), vol. 292, no. 3, pp. 355–363, Mar. 2009, doi: 10.1002/ar.20869.

[34] R. S. Weinstein, R. L. Jilka, A. M. Parfio, and S. C. Manolagas, “Inhibition of osteoblastogenesis and promotion of apoptosis of osteoblasts and osteocytes by glucocorticoids. Potential mechanisms of their deleterious effects on bone,” J Clin Invest, vol. 102, no. 2, pp. 274–282, July 1998, doi: 10.1172/JCI2799.

[35] C. M. Mazur et al., “Partial prevention of glucocorticoid-induced osteocyte deterioration in young male mice with osteocrin gene therapy,” iScience, vol. 25, no. 9, p. 105019, Sept. 2022, doi: 10.1016/j.isci.2022.105019.

[36] M. N. Wein et al., “SIKs control osteocyte responses to parathyroid hormone,” Nat Commun, vol. 7, p. 13176, Oct. 2016, doi: 10.1038/ncomms13176.

[37] A. J. Aguirre et al., “Genomic Copy Number Dictates a Gene-Independent Cell Response to CRISPR/Cas9 Targeting,” Cancer Discov, vol. 6, no. 8, pp. 914–929, Aug. 2016, doi: 10.1158/2159-8290.CD-16-0154.

[38] B. Adamson et al., “A Multiplexed Single-Cell CRISPR Screening Plaqorm Enables Systematic Dissection of the Unfolded Protein Response,” Cell, vol. 167, no. 7, pp. 1867–1882.e21, Dec. 2016, doi: 10.1016/j.cell.2016.11.048.

[39] C. I. De Zeeuw et al., “CLIP-115, a novel brain-specific cytoplasmic linker protein, mediates the localization of dendritic lamellar bodies,” Neuron, vol. 19, no. 6, pp. 1187–1199, Dec. 1997, doi: 10.1016/s0896-6273(00)80411-0.

[40] C. C. Hoogenraad, A. Akhmanova, F. Grosveld, C. I. De Zeeuw, and N. Galjart, “Functional analysis of CLIP-115 and its binding to microtubules,” J Cell Sci, vol. 113 (Pt 12), pp. 2285–2297, June 2000, doi: 10.1242/jcs.113.12.2285.

[41] J. Gao et al., “Endoplasmic reticulum mediates mitochondrial transfer within the osteocyte dendritic network,” Sci. Adv., vol. 5, no. 11, p. eaaw7215, Nov. 2019, doi: 10.1126/sciadv.aaw7215.

[42] J. G. Doench, “Am I ready for CRISPR? A user’s guide to genetic screens,” Nat Rev Genet, vol. 19, no. 2, pp. 67–80, Feb. 2018, doi: 10.1038/nrg.2017.97.

[43] M. Alkobtawi, P. Pla, and A. H. Monsoro-Burq, “BMP signaling is enhanced intracellularly by FHL3 controlling WNT-dependent spatiotemporal emergence of the neural crest,” Cell Rep, vol. 35, no. 12, p. 109289, June 2021, doi: 10.1016/j.celrep.2021.109289.

[44] I. D. Coghill et al., “FHL3 is an actin-binding protein that regulates alpha-actinin-mediated actin bundling: FHL3 localizes to actin stress fibers and enhances cell spreading and stress fiber disassembly,” J Biol Chem, vol. 278, no. 26, pp. 24139–24152, June 2003, doi: 10.1074/jbc.M213259200.

[45] D. Gau and P. Roy, “SRF’ing and SAP’ing - the role of MRTF proteins in cell migration,” J Cell Sci, vol. 131, no. 19, p. jcs218222, Oct. 2018, doi: 10.1242/jcs.218222.

[46] E. H. F. Jandrey et al., “NDRG4 promoter hypermethylation is a mechanistic biomarker associated with metastatic progression in breast cancer patients,” NPJ Breast Cancer, vol. 5, p. 11, 2019, doi: 10.1038/s41523-019-0106-x.

[47] R. H. Zhou, K. Kokame, Y. Tsukamoto, C. Yutani, H. Kato, and T. Miyata, “Characterization of the human NDRG gene family: a newly identified member, NDRG4, is specifically expressed in brain and heart,” Genomics, vol. 73, no. 1, pp. 86–97, Apr. 2001, doi: 10.1006/geno.2000.6496.

[48] J. Chen, K. Yuan, X. Mao, J. M. Miano, H. Wu, and Y. Chen, “Serum response factor regulates bone formation via IGF-1 and Runx2 signals,” J Bone Miner Res, vol. 27, no. 8, pp. 1659–1668, Aug. 2012, doi: 10.1002/jbmr.1607.

[49] C. C. Hoogenraad et al., “Targeted mutation of Cyln2 in the Williams syndrome critical region links CLIP-115 haploinsufficiency to neurodevelopmental abnormalities in mice,” Nat Genet, vol. 32, no. 1, pp. 116–127, Sept. 2002, doi: 10.1038/ng954.

[50] U. Francke, “Williams-Beuren syndrome: genes and mechanisms,” Hum Mol Genet, vol. 8, no. 10, pp. 1947–1954, 1999, doi: 10.1093/hmg/8.10.1947.

[51] C. Schubert, “The genomic basis of the Williams – Beuren syndrome,” Cell. Mol. Life Sci., vol. 66, no. 7, pp. 1178–1197, Apr. 2009, doi: 10.1007/s00018-008-8401-y.

[52] J. M. van Hagen et al., “Contribution of CYLN2 and GTF2IRD1 to neurological and cognitive symptoms in Williams Syndrome,” Neurobiol Dis, vol. 26, no. 1, pp. 112–124, Apr. 2007, doi: 10.1016/j.nbd.2006.12.009.

[53] G. Vandeweyer, N. Van der Aa, E. Reyniers, and R. F. Kooy, “The contribution of CLIP2 haploinsufficiency to the clinical manifestations of the Williams-Beuren syndrome,” Am J Hum Genet, vol. 90, no. 6, pp. 1071–1078, June 2012, doi: 10.1016/j.ajhg.2012.04.020.

[54] H. V. Goodson et al., “CLIP-170 interacts with dynactin complex and the APC-binding protein EB1 by different mechanisms,” Cell MoKl Cytoskeleton, vol. 55, no. 3, pp. 156–173, July 2003, doi: 10.1002/cm.10114.

[55] G. Lansbergen et al., “Conformational changes in CLIP-170 regulate its binding to microtubules and dynactin localization,” J Cell Biol, vol. 166, no. 7, pp. 1003–1014, Sept. 2004, doi: 10.1083/jcb.200402082.

[56] A. Akhmanova et al., “Clasps are CLIP-115 and -170 associating proteins involved in the regional regulation of microtubule dynamics in motile fibroblasts,” Cell, vol. 104, no. 6, pp. 923–935, Mar. 2001, doi: 10.1016/s0092-8674(01)00288-4.

[57] E. J. Lawrence, M. Zanic, and L. M. Rice, “CLASPs at a glance,” J Cell Sci, vol. 133, no. 8, p. jcs243097, Apr. 2020, doi: 10.1242/jcs.243097.

[58] H. Kamioka, Y. Sugawara, T. Honjo, T. Yamashiro, and T. Takano-Yamamoto, “Terminal differentiation of osteoblasts to osteocytes is accompanied by dramatic changes in the distribution of actin-binding proteins,” J Bone Miner Res, vol. 19, no. 3, pp. 471–478, Mar. 2004, doi: 10.1359/JBMR.040128.

[59] R. M. Guerra, V. M. Fowler, and L. Wang, “Osteocyte Dendrites: How Do They Grow, Mature, and Degenerate in Mineralized Bone?,” Cytoskeleton, vol. 82, no. 9, pp. 556–570, Sept. 2025, doi: 10.1002/cm.21964.

[60] C. Mazur, et al., “Trafficking and translation of mRNA in osteocyte dendrites,” Oct. 01, 2024, bioRxiv. doi: 10.1101/2024.10.01.616162.

[61] A. Saidi Brikci-Nigassa et al., “Phosphorylation controls the interaction of the connexin43 C-terminal domain with tubulin and microtubules,” Biochemistry, vol. 51, no. 21, pp. 4331–4342, May 2012, doi: 10.1021/bi201806j.

[62] A. Deb Roy, C. Saez Gonzalez, F. Shahid, E. Yadav, and T. Inoue, “Optogenetically Induced Microtubule Acetylation Unveils the Molecular Dynamics of Actin-Microtubule Crosstalk in Directed Cell Migration,” Dec. 02, 2024, Cell Biology. doi: 10.1101/2024.12.01.626286.

[63] J. P. Kerr et al., “Detyrosinated microtubules modulate mechanotransduction in heart and skeletal muscle,” Nat Commun, vol. 6, p. 8526, Oct. 2015, doi: 10.1038/ncomms9526.

[64] G. Kreitzer, G. Liao, and G. G. Gundersen, “Detyrosination of tubulin regulates the interaction of intermediate filaments with microtubules in vivo via a kinesin-dependent mechanism,” Mol Biol Cell, vol. 10, no. 4, pp. 1105–1118, Apr. 1999, doi: 10.1091/mbc.10.4.1105.

[65] E. Lewkowicz, F. Herit, C. Le Clainche, P. Bourdoncle, F. Perez, and F. Niedergang, “The microtubule-binding protein CLIP-170 coordinates mDia1 and actin reorganization during CR3-mediated phagocytosis,” The Journal of Cell Biology, vol. 183, no. 7, pp. 1287–1298, Dec. 2008, doi: 10.1083/jcb.200807023.

[66] D. J. Mun et al., “Gcap14 is a microtubule plus-end-tracking protein coordinating microtubule–actin crosstalk during neurodevelopment,” Proc. Natl. Acad. Sci. U.S.A., vol. 120, no. 8, p. e2214507120, Feb. 2023, doi: 10.1073/pnas.2214507120.

[67] J. Maurin et al., “The Beta-Tubulin Isotype TUBB6 Controls Microtubule and Actin Dynamics in Osteoclasts,” Front. Cell Dev. Biol., vol. 9, p. 778887, Nov. 2021, doi: 10.3389/fcell.2021.778887.

[68] J. M. Spatz et al., “The Wnt Inhibitor Sclerostin Is Up-regulated by Mechanical Unloading in Osteocytes in Vitro,” J Biol Chem, vol. 290, no. 27, pp. 16744–16758, July 2015, doi: 10.1074/jbc.M114.628313.

[69] A. Dobin et al., “STAR: ultrafast universal RNA-seq aligner,” Bioinformatics, vol. 29, no. 1, pp. 15–21, Jan. 2013, doi: 10.1093/bioinformatics/bts635.

[70] S. Anders, P. T. Pyl, and W. Huber, “HTSeq--a Python framework to work with high-throughput sequencing data,” Bioinformatics, vol. 31, no. 2, pp. 166–169, Jan. 2015, doi: 10.1093/bioinformatics/btu638.

[71] R. Patro, G. Duggal, M. I. Love, R. A. Irizarry, and C. Kingsford, “Salmon provides fast and bias-aware quantification of transcript expression,” Nat Methods, vol. 14, no. 4, pp. 417–419, Apr. 2017, doi: 10.1038/nmeth.4197.

[72] M. I. Love, W. Huber, and S. Anders, “Moderated estimation of fold change and dispersion for RNA-seq data with DESeq2,” Genome Biol, vol. 15, no. 12, p. 550, 2014, doi: 10.1186/s13059-014-0550-8.

[73] K. R. Sanson et al., “Optimized libraries for CRISPR-Cas9 genetic screens with multiple modalities,” Nat Commun, vol. 9, no. 1, p. 5416, Dec. 2018, doi: 10.1038/s41467-018-07901-8.

[74] C. Niger, F. D. Howell, and J. P. Stains, “Interleukin-1beta increases gap junctional communication among synovial fibroblasts via the extracellular-signal-regulated kinase pathway,” Biol Cell, vol. 102, no. 1, pp. 37–49, Oct. 2009, doi: 10.1042/BC20090056.

[75] D. R. Stirling, M. J. Swain-Bowden, A. M. Lucas, A. E. Carpenter, B. A. Cimini, and A. Goodman, “CellProfiler 4: improvements in speed, utility and usability,” BMC Bioinformatics, vol. 22, no. 1, p. 433, Sept. 2021, doi: 10.1186/s12859-021-04344-9.

[76] C. A. Schneider, W. S. Rasband, and K. W. Eliceiri, “NIH Image to ImageJ: 25 years of image analysis,” Nat Methods, vol. 9, no. 7, pp. 671–675, July 2012, doi: 10.1038/nmeth.2089.

